# Integrative pan-cancer analysis reveals a common architecture of dysregulated transcriptional networks characterized by loss of enhancer methylation

**DOI:** 10.1101/2023.09.22.559009

**Authors:** Jørgen Ankill, Zhi Zhao, Xavier Tekpli, Elin H. Kure, Vessela N. Kristensen, Anthony Mathelier, Thomas Fleischer

## Abstract

Aberrant DNA methylation contributes to gene expression deregulation in cancer. However, these alterations’ precise regulatory role and clinical implications are still not fully understood. In this study, we performed expression-methylation Quantitative Trait Loci (emQTL) analysis to identify deregulated cancer-driving transcriptional networks linked to CpG demethylation pan-cancer. By analyzing 33 cancer types from The Cancer Genome Atlas, we identified and confirmed significant correlations between CpG methylation and gene expression (emQTL) in *cis* and *trans*, both across and within cancer types. Bipartite network analysis of the emQTL revealed groups of CpGs and genes related to important biological processes involved in carcinogenesis; specifically, we identified three types of emQTL networks associated with alterations linked to the regulation of proliferation, metabolism, and hormone-signaling. These bipartite communities were characterized by loss of enhancer methylation in transcription factor binding regions (TFBRs) located in enhancers. The underlying CpGs were topologically linked to upregulated genes through chromatin loops. Loss of enhancer methylation and target genes were exemplified in pancreatic cancer. Penalized Cox regression analysis showed a significant prognostic impact of the pan-cancer emQTL. Taken together, our integrative pan-cancer analysis reveals a common architecture of aberrant DNA demethylation that illustrates a convergence of pathological regulatory mechanisms across cancer types.

## Introduction

Alterations in transcriptional programs are determinants of cancer cell phenotypes. Such changes occur during carcinogenesis, in which normal cells transform into cancer cells by accumulating genetic and epigenetic alterations. When such alterations occur in regulatory regions of the genome, they alter transcriptional activity and RNA levels, adversely affecting cellular function and initiating carcinogenesis. Epigenetic alterations such as DNA methylation are frequently observed in cancer and are considered a hallmark of many cancer types [1]. Aberrant DNA methylation patterns are commonly observed early during cancer development and have been linked to clinically relevant features such as tumor stage, prognosis, and response to therapies [1–3].

During cancer progression, oncogenes tend to be activated through promoter hypomethylation while tumor suppressor genes become inactivated through promoter hypermethylation [4–6]. However, many alterations in DNA methylation occur in more distal intergenic regions. Indeed, DNA methylation on distal regulatory regions such as enhancers, is a strong driver of transcriptional regulation. Enhancers are *cis*-acting regulatory regions able to induce gene transcription up to several megabases away from their target gene. To control transcription, enhancers are bound by key proteins such as transcription factors (TFs) and are brought close 3D proximity with their target genes through the formation of chromatin interaction loops [7, 8]. Enhancer hypermethylation is associated with transcriptional inactivity, while enhancer hypomethylation associates with TF binding and transcriptional activation of distal-looped genes [9, 10]. TFs are DNA-binding proteins interacting with specific genomic motifs to regulate gene expression. DNA-binding elements tend to be located in unmethylated *cis*-regulatory regions [11]. While some TFs are expressed by most cells, others are cell-type specific and are critical for cell fate determination [12].

Hypermethylation and loss of TF binding can alter chromatin interactions in cancer [13]. Moreover, DNA methylation can impair TF binding; conversely, specific TFs can promote DNA hypomethylation [14]. For instance, TF binding can hinder DNA methyltransferase (DNMTs) access to DNA. Moreover, pioneer TFs, which can engage with heterochromatin to trigger chromatin opening can interact with histone-modifying proteins and demethylation-competent Ten Eleven Translocation (TETs) enzymes to trigger local DNA demethylation, as demonstrated for FOXA1 and CEBPα [14]. Pioneer factors can also disrupt the activity of DNMTs responsible for maintaining DNA methylation, consequently leading to the loss of DNA methylation during DNA replication and cell division through passive demethylation [14]. Additionally, passive demethylation can occur non-enzymatically when DNA maintenance machinery fails to efficiently re-store the lost methylation pattern after each round of cell division. Furthermore, abnormal acitivty of epigenome gene modifying enzymes can influence the genetic landscape and promote intratumoral genetic heterogeniety [15–17]. The precise impact of changes in DNA methylation patterns on TF binding and, cancer development and progression remain incompletely understood. To comprehend tumor phenotypes and their implication on clinical outcomes, it is crucial to better characterize the crosstalk between the epigenome and the transcriptome. Moreover, identifying commonalities and differences in epigenetically dysregulated transcriptional networks across cancer-types will pave the way to clinical strategies that may benefit patients with different types of cancer.

We previously computed expression-methylation quantitative trait loci (emQTL) to reveal associations between enhancer methylation and estrogen signaling and ER-independent proliferation in breast cancer [18, 19]. These studies highlighted that epigenetic features of enhancer regions drove the regulation of gene expression, the activity of TFs at these regions, and chromatin loops [18, 19]. Additionally, we identified emQTL links that derive from the influence of non-cancer cell infiltration on bulk tumor measurements of DNA methylation and gene expression; the identified associations reflected varying immune cell and fibroblast infiltration. To comprehend how DNA methylation associates with gene expression on a genome-wide scale in multiple cancers and its clinical implications, we performed genome-wide emQTL analysis pan-cancer to identify disease-driving epigenetically dysregulated transcriptional networks linked to loss of DNA methylation. By integrating DNA methylation and gene expression data from the 33 different cancer types in TCGA, we identified a cell cycle-related community of emQTL characterized by the loss of enhancer methylation situated in transcription factor binding regions for FOSL1/2 and JUN (parts of the AP-1 complex) with concomitant upregulation of proliferation-related genes. The proliferation-related CpGs and genes were functionally connected through predicted chromatin loops. Interestingly, loss of enhancer methylation and upregulation of the proliferation-related genes were observed in most cancer types, suggesting that proliferation is epigenetically regulated in several cancer types. Two novel cancer-related communities of emQTL were discovered and linked to the regulation of metabolism and hormone-signaling in cancer. Another community of emQTL was found linked to the tumor microenvironment, highlighting the link between tumor-infiltrating cells and the cancer cell epigenome. Our findings provide insights into the regulatory mechanisms of the cancer epigenome on the transcriptome and identify potential targets for clinical intervention.

## Results

### Identification of pan-cancer expression-methylation quantitative trait loci (emQTL)

To identify robust associations between DNA methylation and gene expression, in *cis* and *trans*, we correlated genome-wide levels of DNA methylation and gene expression from the TCGA-PANCAN dataset (see Material and Methods; Supplementary figure 1 for workflow outline). The TCGA pan-cancer dataset was split into a discovery (n=4104) and a validation cohort (n=4089; see Material and methods). 1 192 567 (87%) out of the 1 363 565 significant CpG-gene associations discovered were significant in the validation cohort. Most correlations had negative Pearson coefficient values (64.8% negative vs 35.2% positive; Supplementary figure 2A-C), indicating that methylation at a CpG increases when the level of expression for the correlated gene decreases, which can reveal a potentially putative inhibitory effect of CpG methylation on gene expression. The validated emQTL includes 5472 genes and 45 406 CpGs. We found *cis*-associations (same chromosome; 83 035 associations) to be significantly enriched compared to *trans* associations (1 037 323 associations; hypergeometric test p-value<2.2e-16; fold enrichment (FE)=1.42). The validated associations are hereafter called emQTL. Although causality on gene expression regulation cannot be determined from statistical correlations, the emQTL provides insight into the overall degree of coordination between CpG methylation and gene expression globally, considering both potential direct and indirect regulation.

### Characterization of the pan-cancer emQTL by bipartite network analysis reveals communities with distinct biology

To identify coordinated emQTL, we performed bipartite network analysis using Complex Network Descriptions Of Regulators (CONDOR) [20], identifying communities of highly connected emQTL-CpGs and emQTL-genes. Since our study focused on the events where DNA methylation had a putative inhibitory effect on gene expression, we considered negative correlations in the community detection analysis (772 730 emQTL). We detected six pan-cancer emQTL communities (Figure 1A-B; Supp. table 1A-B). To elucidate the biological functions of the emQTL communities, we performed gene set enrichment analyses (GSEA) using gene sets obtained from the Molecular Signatures Database (MSigDB) [21]. We found the communities to be enriched for genes involved in metabolic processes (Community 1), cell cycle (Community 2), neuroendocrine-like (NE-like) functions (Community 3), hormone-signaling (Community 4), immune response (Community 5) and epithelial-mesenchymal transition (EMT; Community 6; Figure 1C, Supp. table 1C). We confirmed the consistency of the negative correlations between emQTL-CpGs and emQTL genes in each cancer type (Figure 1D). The consistency of most communities suggests that the emQTL represents both intra- and inter-cancer type features.

**Figure 1.**
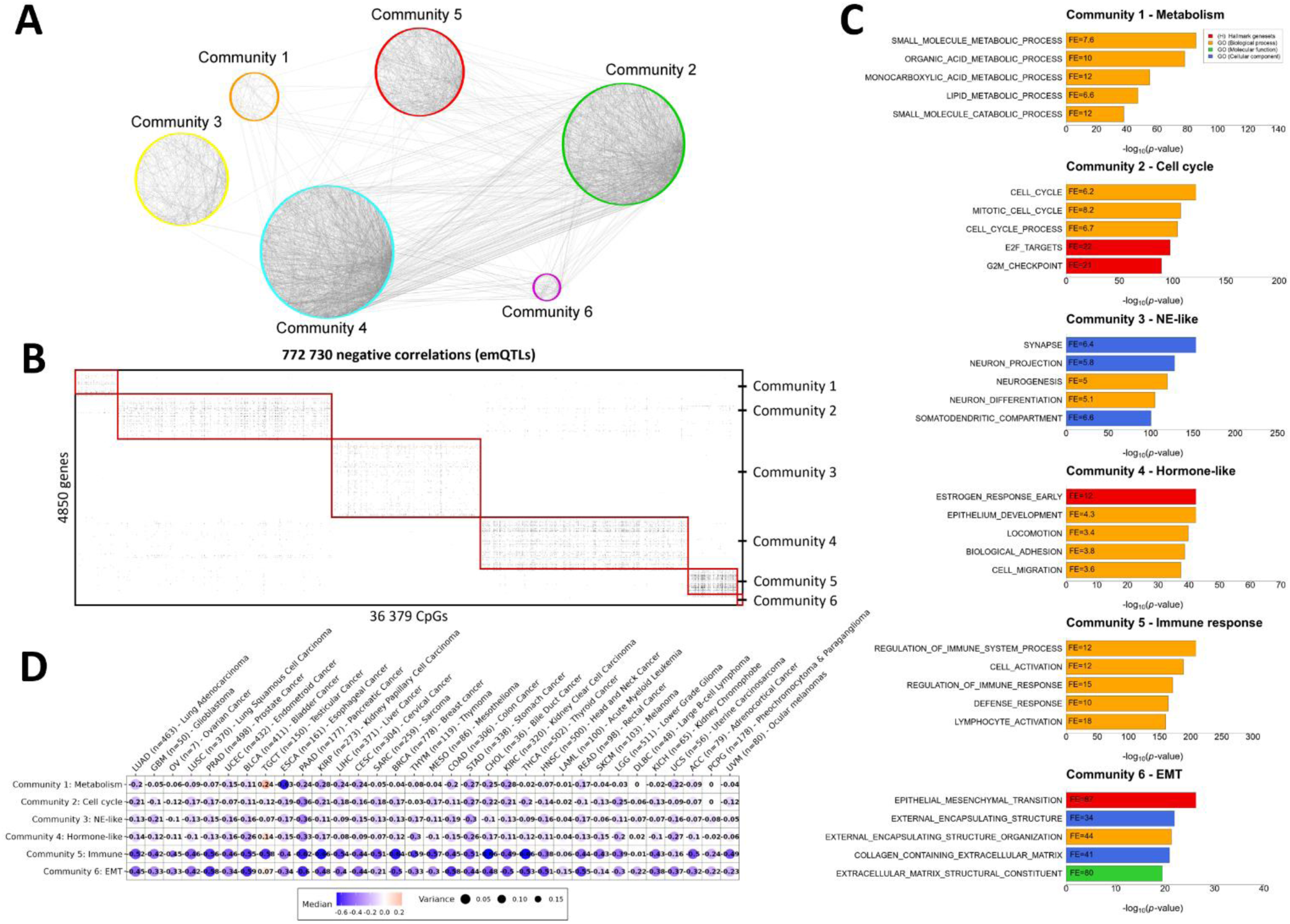
Characterization of the pan-cancer emQTL. (A) Interaction plot showing the emQTL associations using Cytoscape v.3.9.1 [22]. For clarity of visualization, only 5000 intra- and inter-community associations are shown (4808 randomly selected emQTL related to communities 1 to 5 and all 192 emQTL related to community 6). (B) Heatmap showing the pan-cancer emQTL found in the TCGA-PANCAN dataset (772 730 emQTL) sorted by emQTL community in which black points indicate significant CpG-gene associations (emQTL). Rows represent genes and the columns represent CpGs involved in emQTL. (C) GSEA of the genes in communities 1 to 6 using the Hallmark and gene ontology C5 (GO biological process, GO molecular function, and GO cellular component) gene set collections from the MSigDB [23]. The bar length represents the −log10-transformed Benjamini-Hochberg (BH) corrected p-value obtained by hypergeometric testing. Only the top 5 most significantly enriched gene sets for each community are shown. (D) Dot plot showing the median Pearson coefficient obtained by reanalyzing the pan-cancer emQTL in each community for the 33 cancer types separately. The color of the dot represents the sign of the median correlation coefficients (i.e., median negative correlation coefficient in blue and median positive correlation coefficient in red). Dot size denotes the variance of the emQTL correlation coefficients.

### The emQTL-CpGs reside *in cis* regulatory regions

To assess the functional context of the pan-cancer emQTL-CpGs, we assessed their enrichment in specific categories of regulatory regions by considering the pan-tissue candidate *cis*-Regulatory Elements (cCREs) predicted by ENCODE [24]. We found the emQTL-CpGs from all communities significantly enriched in enhancer regions (Figure 2A, Supp. table 1D). Moreover, the CpGs in communities 1 to 5 were enriched in promoter regions. Since TFs can bind to promoter and enhancer regions to modulate gene expression, we aimed to identify which TFs may bind to these regions using TF-DNA interaction data obtained from the UniBind database [25]. We found the emQTL-CpGs in each community to be enriched in TF binding regions (TFBR) of TFs associated with functions coherent with the GSEA of the communities (Figure 2B, Supp. table 1E). Supplementary figure 4 shows the overlap between the TFBR for the top enriched TFs from each community.

**Figure 2.**
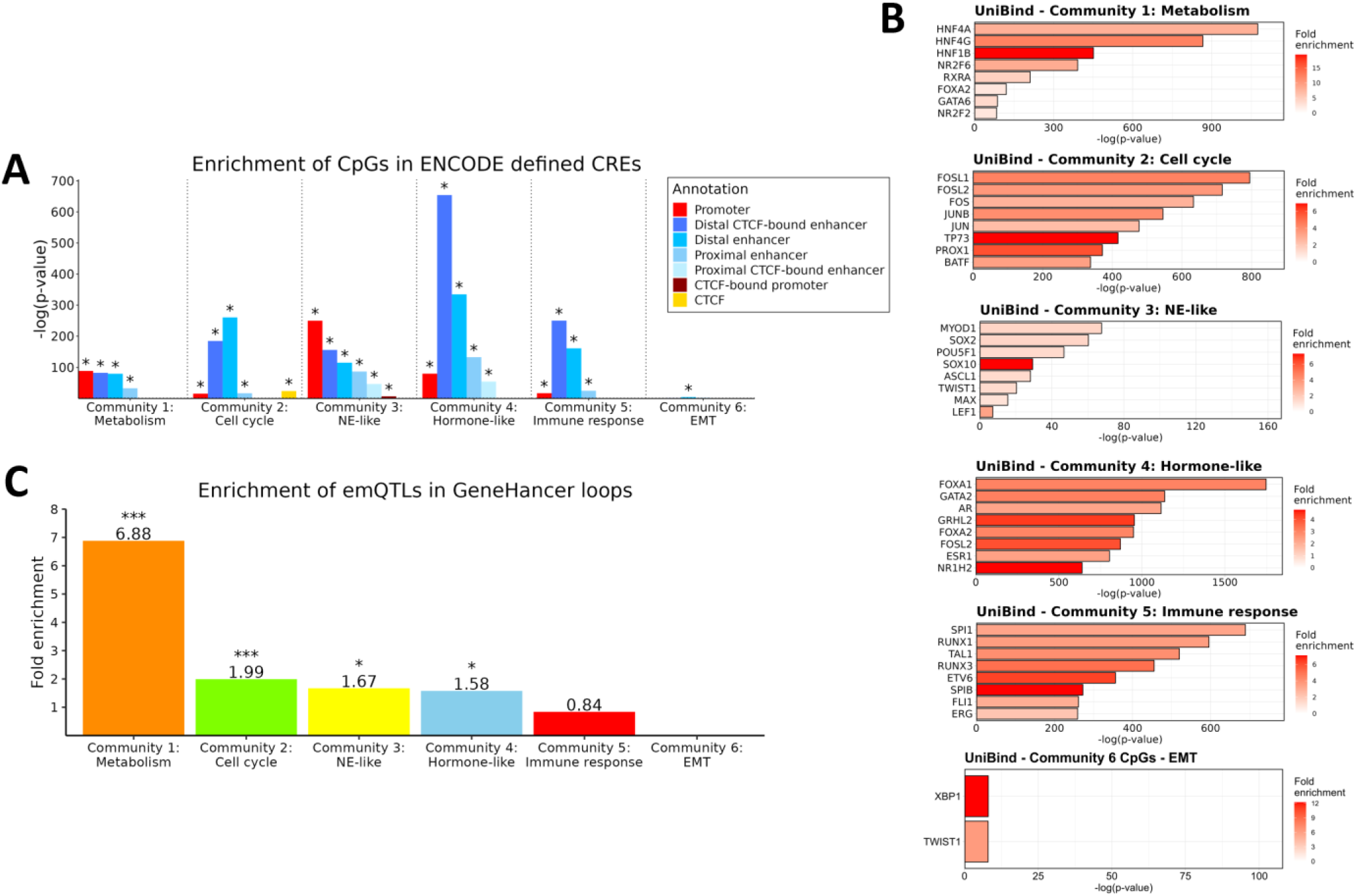
Genomic location of the pan-cancer emQTL. (A) Enrichment of emQTL-CpGs in ENCODE-defined cCREs. The height of the bars represents the log-transformed BH-corrected p-value obtained by hypergeometric testing using the 450k probes as background. Significant enrichments (BH-corrected p-value<0.05) are marked with an asterisk. (B) Enrichment of emQTL-CpGs in TFBR by emQTL community according to UniBind [25]. The length of the bars corresponds to the log-transformed p-values obtained by hypergeometric testing. Only the top most significantly enriched TFs are shown in each plot. (C) Bar plot showing the enrichment of emQTL from each emQTL community in GeneHancer loops. The height of the bars represents the fold enrichment measured as the ratio between the frequency of emQTL found in the head and tail of GeneHancer loops over the expected frequency is such overlaps were to occur at random. Statistically significant enrichments BH-correction obtained by hypergeometric test are marked with an asterisk (* is p<0.05, ** is p<0.001, *** is p<0.0001). See Materials and Methods for a detailed description of the calculation of enrichment.

Enhancers can regulate gene expression through the binding of TFs and the formation of chromatin loop interactions with their target gene. As all emQTL communities were enriched for enhancer regions, we assessed whether the emQTL-CpGs and genes were linked through predicted loops from the GeneHancer [44]. Our analysis revealed that the emQTL in communities 1 to 4 were significantly enriched in GeneHancer loops (Figure 2C, Hypergeometric test p-value=5.50e-11, 2.74e-05, 3.59e-03, 7.97e-04 respectively; Supp. table 1F). No statistically significant enrichment in loops was observed for the emQTL in the immune and EMT communities. The results from the emQTL communities are summarized in Table 1.

**Table 1.** Summary of the pan-cancer emQTL communities.

| Community | Number of CpGs | Number of genes | GSEA function (MSigDB) | Enriched ENCODE-defined cCRE | Enriched UniBind TFs | Enrichment in GeneHancer associations |
| --- | --- | --- | --- | --- | --- | --- |
| 1 | 2207 | 447 | Metabolism | Promoters, enhancers | HNF4A/G/B and RXRA [26-29] | Enriched |
| 2 | 11 517 | 892 | Cell cycle | Promoters, enhancers, CTCF binding sites | FOSL1/2 and JUN [30-32] | Enriched |
| 3 | 8136 | 1560 | NE-like | Promoters, enhancers | MYOD1, SOX2, POU5F1 [33-35] | Enriched |
| 4 | 11 166 | 1067 | Hormone-like | Promoters, enhancers | ESR1 [36] and AR [37] | Enriched |
| 5 | 2674 | 529 | Immune response | Promoters, enhancers | SPI1 [38], RUNX1/3 [39, 40] and TAL1 [41] | Not significant |
| 6 | 31 | 43 | EMT | Enhancers | TWIST1 and XBP1 [42, 43] | Not significant |

### The pan-cancer emQTL can predict survival in many cancer types

To assess the ability of the pan-cancer emQTL to predict prognosis, we built Cox regression with a ridge penalty to contract models for survival predictions by estimating the hazard function (Ridge Cox model) [45]. Specifically, we performed principal component analysis (PCA) on the z-scores obtained from methylation and expression data for the emQTL-CpGs and genes to standardize the data on the same scale (see Material and Methods. In each community, the first two principal components (PCs) of emQTL-CpGs and -genes were extracted. We regressed patients’ survival in each cancer type by Ridge Cox model and obtained Uno’s C-index. For comparison, we extracted the first two PCs from the emQTL-CpGs and emQTL-genes respectively, i.e., without considering emQTL communities, and regressed patients’ survival in each cancer type on the four PCs from all the emQTL communities by Ridge Cox model. Uno’s C index was estimated to evaluate the models in which a value below 0.5 indicates a poor model and a value above 0.7 indicates a good model. The PCs from the pan-cancer emQTL considering the six communities show the predictive power of patients’ survival (Figure 3A), and had significantly higher C-indexes than the four PCs from the pan-cancer emQTL without considering the communities (p-value=1.41e-11; Supplementary figure 3). Figure 3B-E shows examples of differences in survival outcomes between the predicted high and low-risk group for the ocular melanoma, adrenocortical cancer, mesothelioma, and lower-grade glioma (C-index=0.81, 0.75, 0.75 and 0.68 respectively). Altogether our analysis highlights the ability of the pan-cancer emQTL to predict survival outcome. We hypothesize that the emQTL has an important role in cancer-related regulation of transcriptional networks.

**Figure 3.**
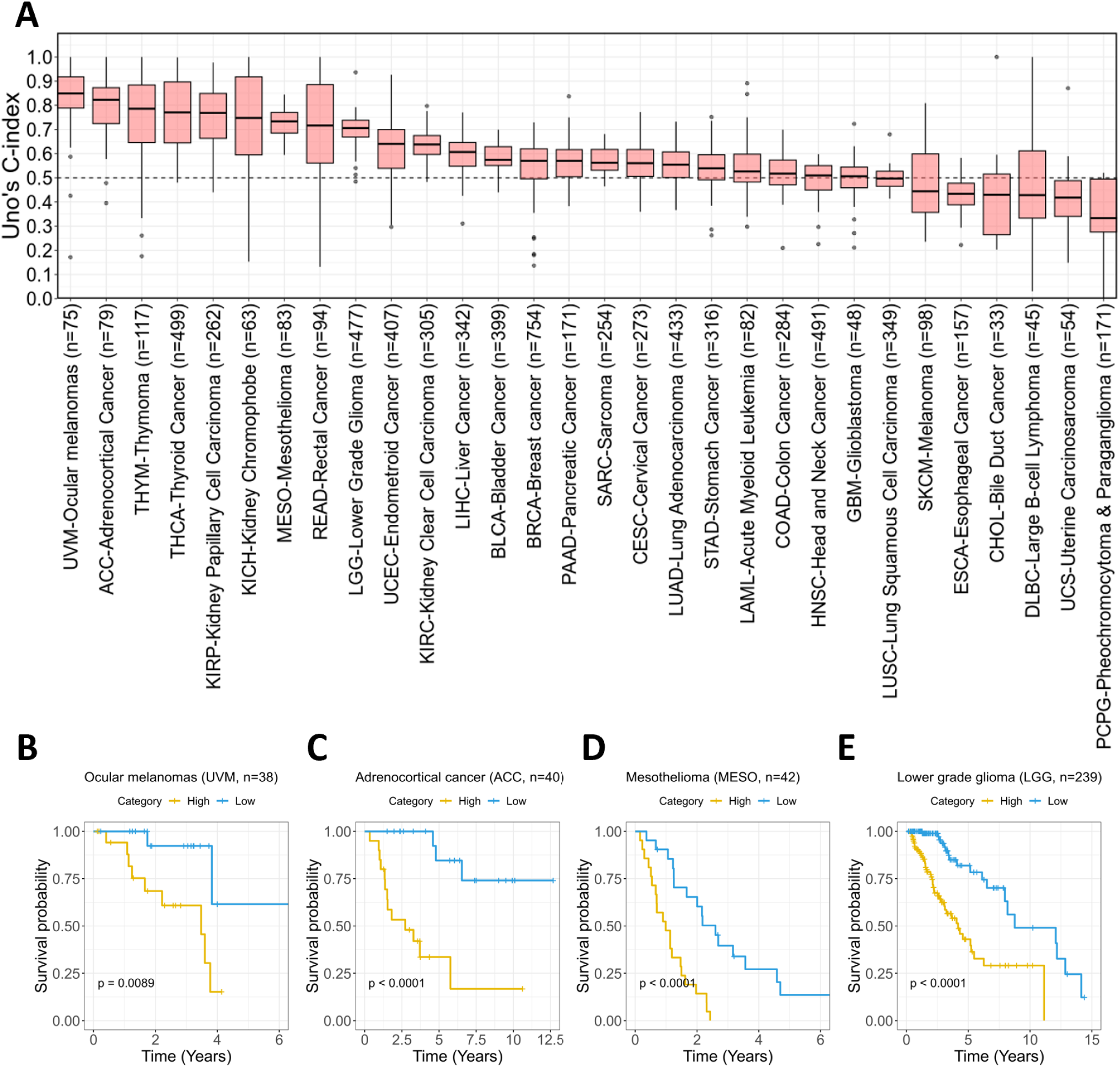
Survival prediction of the pan-cancer emQTL. (A) The pan-cancer patients were randomly split into 80% training and 20% test data 50 times. Each Box shows Uno’s C-indexes estimated from the randomly split 20% test data. Due to few samples for ovarian cancer (n=8) and high censoring in prostate- and testicular cancer, these cancer types were not included in the analysis. Statistically significant differences in C-indexes between the with and without community groups for each cancer type after BH-correction are denoted with an asterisk (* is p<0.05, ** is p<0.001, *** is p<0.0001). Kaplan-Meier survival curves and log-rank tests show the ability of the pan-cancer emQTL to discriminate patients into high- and low-risk groups in regards to survival in ocular melanoma (B), adrenocortical cancer (C), mesothelioma (D) and lower grade glioma (E). Patients were divided into two categories based on the risk scores obtained from the Cox models with ridge penalty. The pan-cancer patients for each cancer type were randomly split into 50% training and 50% test data.

### The cell cycle community highlights important processes of cancer pathogenesis

To shed light on the pathogenic molecular processes associated with the cell cycle community, we performed hierarchical clustering of DNA methylation levels at CpGs and expression of genes in the cell cycle community using the TCGA-PANCAN dataset. We found the methylation and expression profiles to reflect the tissue of origin of the tumors (Figure 4A and D; Chi-square test p-value<2.2e-16 (three patient subclusters)) which is consistent with previous reports [46]. Moreover, we observed that the cell cycle community CpGs are hypomethylated in most cancer types compared to normal adjacent tissue (Figure 4B). The hypomethylation is specifically observed at the TFBRs of FOSL1/2 and JUN (Supplementary figure 5A). We further assessed whether the CpGs and genes associated with the cell cycle community were located in open chromatin regions using ATAC-seq data [47]. As expected, the cancer types harboring the open chromatin around the cell cycle community CpGs were also the cancer types with the lowest level of DNA methylation at these sites (Figure 4C). Concomitantly, the genes associated with the cell cycle community show significantly higher expression in tumor cells than in normal cells for most cancer types (Figure 4E). Finally, the cancer types with the highest expression level of the cell cycle community genes correspond to the ones with the highest open chromatin signal (Figure 4F). These lines of evidence show that the cell cycle community CpGs are hypomethylated in most cancer types and are found in accessible enhancer regions forming interactions with transcription start site (TSS) within open chromatin regions regulating cell cycle-related genes.

**Figure 4.**
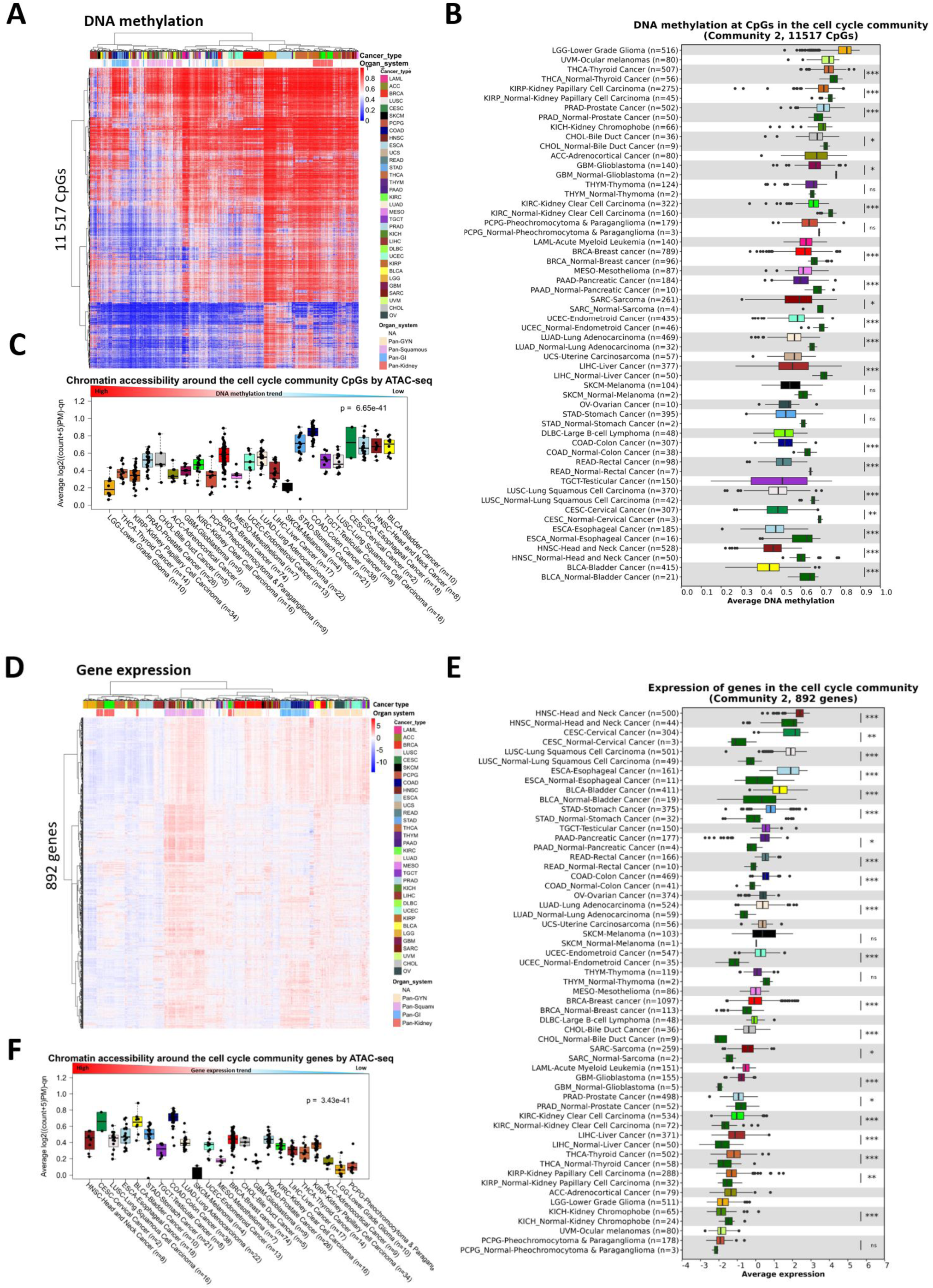
DNA methylation at CpGs and expression of genes in the cell cycle community. (A) Hierarchical clustering of DNA methylation levels at CpGs in the cell cycle community in the TCGA-PANCAN dataset (n=8229). Rows represent CpGs and columns represent tumor samples. Unmethylated CpGs are shown in blue and methylated CpGs are in red. Histopathological features including cancer types and organ system are indicated in the columns and CpGs in TFBRs of FOSL1/2 and JUN is annotated in the rows. (B) DNA methylation at the cell cycle community CpGs by cancer type in the TCGA-PANCAN dataset. Normal samples are included for those cancer types with data available from TCGA. BH-corrected Wilcoxon-rank sum test p-values are denoted. (C) Box plot showing the chromatin accessibility (based on ATAC-seq) in the region at the cell cycle community CpGs obtained from the TCGA-PANCAN dataset. A high value indicates open chromatin while a low value indicates more compact chromatin. The cancer types are ordered by the median DNA methylation levels at the CpGs in the cell cycle community. (D) Unsupervised hierarchical clustering of gene expression levels of the cell cycle community genes in TCGA-PANCAN dataset (n=9875). Genes are found in the rows and the tumor samples are in the columns. Red indicates high expression and blue indicates low expression. Annotations of cancer type and organ system are included. (E) Average expression of the cell cycle community genes by cancer type. As a comparison, expression values from normal adjacent tissues are included if available for the specific cancer type. BH-corrected Wilcoxon rank-sum test p-values are denoted. (F) shows the chromatin accessibility around the cell cycle community genes in the TCGA-PANCAN dataset obtained by ATAC-seq. The cancer types are ordered by the median expression levels of the genes in the cell cycle community for each particular cancer type.

### The hormone-like and metabolism community links alteration in enhancer methylation to variations in gene expression through looping

Hierarchical clustering of DNA methylation levels at the hormone-like and metabolism community CpGs separated the tumors based on the tissue of origin, but also indicated variability in DNA methylation within each cancer type (Supplementary figure 6A-B). The CpGs in both communities were significantly enriched in enhancer regions and were less methylated in most cancer types compared to normal adjacent tissue (Supplementary figure 6C-D). Loss of methylation was also observed specifically at the TFBR of FOXA1, ESR1, and AR at the CpGs in the hormone-like community and at the HNF4A/G and RXRA TFBR in the metabolism community (Supplementary figure 5B-C). By assessing the chromatin accessibility around the hormone-like and metabolism community CpGs, we found lower DNA methylation levels at the CpGs to be associated with more accessible chromatin (Supplementary figure 6E-F). Similarly, to the methylation data, the expression of genes in the hormone-like and metabolism-related communities clustered according to tissue of origin, and variability in expression withing the cancer types was also observed (Supplementary figure 7A-B). We next compared the expression of the hormone-like and metabolism community genes between tumor and normal tissue and found genes in both communities to be significantly higher expressed in tumor tissue compared to normal adjacent tissue for most cancer types (Supplementary figure 7C-D). Low DNA methylation at the hormone-related and metabolism-related CpGs was associated with accessible chromatin, while high expression of the genes was associated with more accessible chromatin (Supplementary figure 7E-F). The emQTL in both communities were ignorantly enriched in inferred GeneHancer enhancer-promoter loops, thereby suggesting a possible functional regulation of hormone-related and metabolism genes. Altogether, our findings suggest that enhancer methylation is linked to the upregulation of hormone- and metabolism genes through chromatin looping in across cancer types.

Adipocytes are a prominent cell type of the tumor microenvironment in solid tumors and express high levels of metabolic genes involved in energy storage and mobilization. To ensure the signal form the metabolism community was not caused by infiltration of non-tumor cells such as adipocytes, we correlated the average DNA methylation levels at the CpGs genes in the metabolism community with the ASCAT tumor purity estimates [48]. No statistically significant correlation was observed between methylation and tumor purity in TCGA (Supplementary figure 9, p-value=0.90). A significant correlation was observed between the average expression of the metabolism genes and tumor purity (p-value=8.54e-06, r-value=−0.05); however, the Pearson correlation coefficient r was close to zero, suggesting that the variations in methylation and expression profiles of the metabolism community CpGs and genes are caused by differences between the tumor cells themselves.

### The immune community reflects variations in the tumor microenvironment

Unlike the DNA methylation levels at the CpGs and expression levels at the genes in the cell cycle, metabolism, and hormone-like communities, CpGs and genes in the immune community did not segregate the tumors equally well according to cancer type (Supplementary figure 10A-D). Since immune cells are prominent in the tumor microenvironment and have different DNA methylation and gene expression profiles compared to cancer cells, we hypothesize that this community of emQTL was influenced by and reflects variation in immune cell infiltration in tumors. To address this, we used the xCell deconvolution [49] algorithm to quantify the level of immune infiltration using gene expression data from TCGA. We divided the tumors into quartile groups based on the level of immune infiltration. We observed a significant difference in DNA methylation levels at the CpGs in the immune community between the quartile groups in which low methylation at CpGs in the immune-community was associated with high immune infiltration and high DNA methylation with low infiltration (Supplementary figure 11A, p-value<2.2e-16). Furthermore, DNA methylation levels at the CpGs in the immune community obtained from isolated immune cells, such as leukocytes, monocytes, T-cells, and B-cells, reflected similar methylation levels at the CpGs as the tumors with high immune infiltration. In contrast, cancer cell lines from the National Cancer Institute (NCI-60) Human Tumor Cell Lines Screen showed similar methylation levels as the quartile group with the lowest infiltration. One exception was leukemia cell lines which is reasonable as the cell of origin for this cancer type is lymphocyte and therefore shares similarities with the infiltrating immune cells. We also observed a positive association between immune infiltration and expression of genes in the immune community, i.e., high expression of the immune community genes is linked to high immune infiltration and vice versa (Supplementary figure 11B). To further confirm the link between immune infiltration and the immune community, we obtained copy-number-derived tumor purity estimates from ASCAT [48] and assessed the link between the xCell-derived immune score and tumor purity. We found a negative association between tumor purity and immune cell infiltration, i.e., low tumor purity is associated with high infiltration of immune cells, and vice versa (Supplementary figure 11C). Altogether, these results suggest that the emQTL in the immune community is caused by variations in immune cell infiltration.

### The EMT community reflects varying fibroblast infiltration in tumors

The smallest emQTL community was the EMT community consisting of 31 CpGs and 43 genes representing 120 emQTL. Similar to the immune community, DNA methylation at the CpGs and expression of genes did not segregate the tumors according to cancer types compared to the cell cycle, metabolism, and hormone-like communities (Supplementary figure 12A-D). Since fibroblasts are a prominent mesenchymal cell type of the tumor microenvironment of many solid tumors and carry out functions such as migration and remodeling of the extracellular matrix (ECM) [50], we assessed whether the EMT community could arise from heterogeneity in tumor composition in regards to fibroblast infiltration. We employed the xCell deconvolution tool [49] to estimate the relative quantity of fibroblast infiltration in each tumor sample using gene expression. By dividing the tumor samples into quartile groups based on the severity of fibroblast infiltration, we found that expression of the EMT community genes was positively associated with fibroblast infiltration (Supplementary figure 13A, p<2.2e-16), i.e. high expression of EMT community genes is linked to high fibroblast infiltration. We also observed a significant negative association between DNA methylation and fibroblast infiltration (Supplementary figure 13B, p-value<2.2e-16). Furthermore, the DNA methylation levels of the EMT community CpGs in tumors with severe fibroblast infiltration showed similar methylation profiles as purified fibroblasts. Conversely, all tumor cell lines from the NCI-60 panel showed methylation levels similar to the low infiltration group. To further confirm that the EMT community was linked to fibroblast infiltration, we assessed whether the quartile groups based on the severity of fibroblast infiltration were linked to the ASCAT tumor purity estimates. A significant and negative association was observed between fibroblast infiltration and tumor purity, i.e., high fibroblast infiltration is linked to low tumor purity, and vice versa (Supplementary figure 13C). Taken together, our findings suggest that the EMT community of emQTL is linked to heterogeneity in the severity of fibroblast infiltration in tumors.

### The NE-like community reflects differences in the tissue of origin and heterogeneity in the tumor microenvironment

To study the differences in terms of methylation and expression of the NE-like community CpGs and genes, we performed unsupervised hierarchical clustering of DNA methylation and gene expression levels of the CpGs and genes in the NE-like community. We found the DNA methylation and expression profiles of the CpGs and genes in the NE-like community to cluster by cancer type (Supplementary figure 14A-B). Among the cancer types, the neuron-related cancer types such as lower-grade glioma (LGG), pheochromocytoma & paraganglioma (PCPG), and glioblastomas (GBM) had pronouncedly different DNA methylation and gene expression profiles compared to the other cancer types. These cancer types showed the highest expression of the NE-like community genes and the lowest DNA methylation at the CpGs in the NE-like community (Supplementary figure 14C-D). Moreover, these cancer types had less compact chromatin around the NE-like community CpGs and genes than other cancer types (Supplementary figure 14E-F).

Altogether, our findings suggest that the NE-like community is associated with the cell of origin. The brain-related cancer types have upregulated neural-related genes through low DNA methylation at enhancers. However, this cannot fully explain the community as there are associations between DNA methylation at the CpGs and expression of the genes in the NE-like community within non-brain related cancer types (Figure 1D). The methylation and expression levels differ between tumor and normal adjacent tissue for most cancer types (Supplementary figure 14C-D). To assess whether this could be caused by variations in the tumor microenvironment, we obtained the tumor microenvironment score from xCell [49]. We assessed the link between DNA methylation of CpGs and the expression of genes in the NE-like community to the severity of infiltration. Due to the prominent differences in methylation and expression profiles between the neuron-like cancer types (LGG, PCPG, and GBM) and the other cancer types, we assessed the link between methylation, expression, and tumor infiltration in the neuron-like (NL) and non-neuron-like (NNL) tumors independently. We found the expression of the NE-like community genes to be positively associated with the microenvironment score from xCell in the NNL tumors (Supplementary figure 15A; p-value=1.29e-121). Moreover, a negative association between the tumor microenvironment score and DNA methylation was observed for the NNL, and the methylation levels from the NCI-60 cancer cell lines reflected the methylation levels in the low infiltration group (Supplementary figure 15B; p-value=3.58e-110). A negative association was observed between the tumor microenvironment score and tumor purity (Supplementary figure 15C). Altogether our findings suggest that the variability in DNA methylation at the CpGs and expression of genes in the NE-like community within the cancer types are caused by variations in the cell composition of the tumor microenvironment in the NNL cancer types. For the NL tumors, there was a negative association between the tumor microenvironment score and expression of the genes in the NE-like community (Supplementary figure 15D, p-value=3.31e-70) and a positive association with the level of DNA methylation (Supplementary figure 15E). A negative association between the tumor microenvironment score and tumor purity was also observed and the main difference in purity was between the tumors with severe infiltration and low, moderate, and high infiltration (Supplementary figure 15F). Altogether this suggests that the NE-like community is mainly caused by differences in the tissue of origin between the tumors and that differences in tumor composition cause the variability in expression and DNA methylation within the cancer-types.

### Identification of cancer-driving emQTL

Since emQTL can be both in *cis* and *trans*, and some might rather be passenger emQTL rather than cancer-driving emQTL, we sought to perform emQTL analysis in a knowledge-guided manner to identify candidate emQTL facilitating cancer-promoting functions through direct regulation. By knowing the characteristics of the transcriptional networks linked to the regulation of the cancer-related emQTL communities (i.e., the hormone-like, cell cycle, and metabolism community), we can perform a knowledge-guided analysis by applying several criteria to the selection of emQTL based on the characteristics of the communities (see methods). Using the knowledge-guided emQTL approach, we identify 1770 proliferation-promoting, 1762 metabolism-promoting, and 1062 hormone-signaling-promoting candidate emQTL in cancer (Supp. table 1G). After identifying candidate emQTL we assessed the link between DNA methylation at the candidate emQTL-CpGs and expression of their linked emQTL-gene for each cancer type and found most candidate emQTL to display correlations within several cancer types (Supp. table 1G). One example of a proliferation-promoting candidate emQTL involves cg23320499 and CAMP Responsive Element Binding Protein Like 2 (CREBL2) found in the GO cell cycle gene set (Figure 5A). CREBL2 associates with the emQTL-CpG cg23320499 and is located in a predicted enhancer in the binding site of JUND and CEBPB. Moreover, DNA methylation at cg23320499 was negatively correlated with the expression of CREBL2 in several cancer types such as thymoma (p-value=8.71e-11, r-value=−0.6) and bile duct cancer (Figure 5B; p-value=1.54e-02, r-value=−0.5). In bile duct tumors, a significantly lower methylation level at cg23320499 and higher expression of CRBEL2 were observed compared to normal adjacent tissue (Figure 5C and D, respectively). These results suggest that we can identify promising candidate proliferation-promoting emQTL using the knowledge-guided emQTL approach.

**Figure 5.**
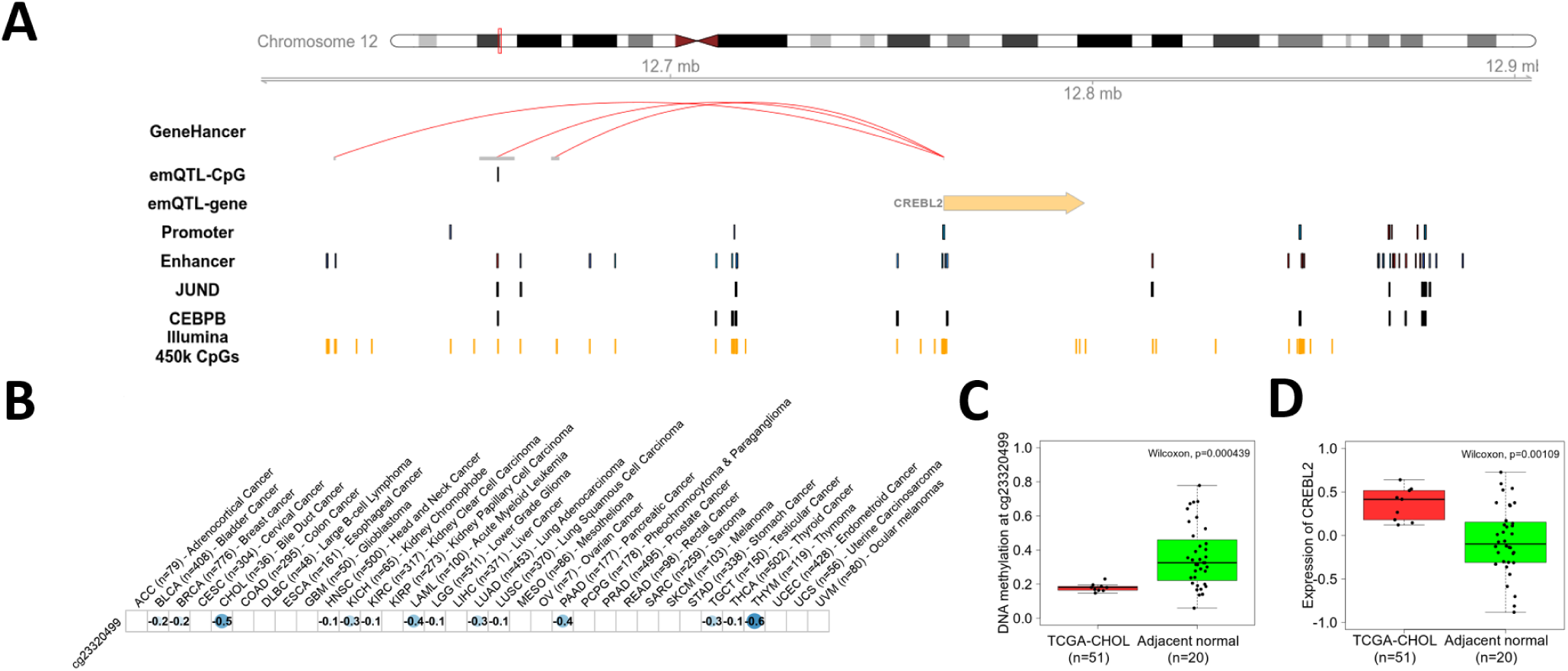
Identification of candidate drivers of proliferative signaling in cancer through knowledge-guided emQTL analysis. (A) Example of a cancer-promoting emQTL identified by knowledge-guided emQTL. The emQTL-CpG (cg23320499) is found in an Encode-defined enhancer region in the binding region of JUND and CEBPB and can form interactions with the CREBL2 promoter. (B) Correlation between DNA methylation at cg23320499 and expression of CREBL2 in each of the 33 cancer types in the TCGA-PANCAN dataset. A blue color indicates a negative correlation and a red color indicates a positive correlation. The size of the dot represents the strength of the correlation. The Pearson correlation coefficient r is denoted in each box of the dot plot. Only significant correlations after BH-correction are represented with a dot. Box plots showing the difference in DNA methylation at cg23320499 (C) and expression of CREBL2 (D) in bile duct cancer (CHOL) and normal adjacent tissue. Wilcoxon test p-values are denoted.

### Assessing the role of the cancer-related emQTL in different cancer types

Next, we sought to study the role of the pan-cancer emQTL of the cancer-related communities (i.e., cell cycle, metabolism, and hormone-like communities) in further detail in a cancer-specific manner as case examples to show the link between DNA methylation at the emQTL-CpGs and target gene expression.

#### The cell cycle community

Among the cancer types, pancreatic tumors exhibited the strongest negative correlation between methylation at the CpGs and expression of the genes in the cell cycle community, as demonstrated in Figure 1D. Due to the aggressive nature of these tumors and the limited treatment options available, we chose to deepen our understanding of the disease by further investigating the role of the pan-cancer emQTLs in the cell cycle community in this particular cancer type. By clustering the DNA methylation levels at the CpGs in the cell cycle community in pancreatic cancer tumors from TCGA, we found a prominent variability in DNA methylation profiles across the tumors (Figure 6A). The CpGs were significantly less methylated in pancreatic cancer tumors compared to normal adjacent tissue both in the TCGA dataset and in the independent cohort from pancreatic ductal adenocarcinomas (PDAC) obtained from Nones et al. (Figure 6B-C; p-value=2.16e-05 and 6.43e-15 respectively) [51]. To investigate the functional role of the cell cycle community CpGs in pancreatic cancer specifically, we obtained genome annotation data from Segway derived from multiple ChIP-seq experiments of histone modifications from the pancreatic cancer cell line PANC-1 [52, 53]. Our analysis revealed a statistically significant enrichment of the CpGs in enhancer and quiescent regions (p-value=1.77e-80 and 2.31e-76 respectively; Figure 6D; Supp. table 1H). We next performed hierarchical clustering of the gene expression profiles of the cell cycle community genes and observed variability between tumors, with some tumors showing consistently higher expression of these genes (Figure 6E). The cell cycle community genes were significantly more expressed in pancreatic tumors compared to normal tissue obtained from TCGA and an independent cohort with expression data obtained from Sandhu et al. [54] (Figure 6F and G respectively). Having in mind that the emQTL in the cell cycle community was enriched in interaction loops in the pancreatic cancer cell line PANC-1 (p-value=0.0189; Supplementary figure 8; Supp. table 1F), we assessed whether there was a significant correlation between DNA methylation at the CpGs and expression of the genes in loops in pancreatic cancer. A statistically significant correlation was observed (Figure 6I; p-value=2.75e-31, r=−0.73). These findings suggest that loss of enhancer methylation at TFBRs of FOSL1/2 and JUNB is linked to the upregulation of proliferation-related genes in several cancer types including pancreatic cancer.

**Figure 6.**
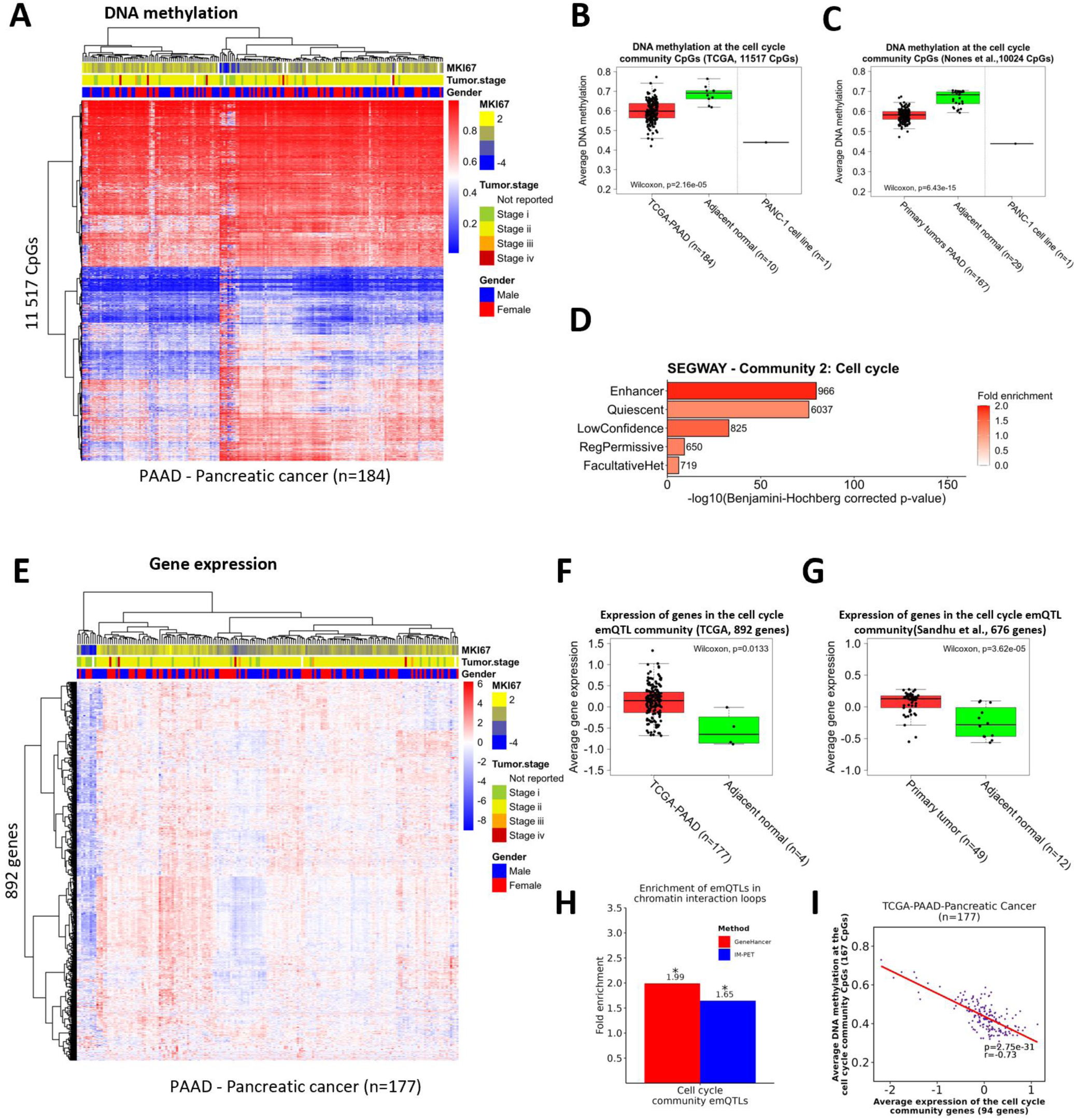
The cell cycle community in pancreatic cancer. (A) Unsupervised hierarchical clustering of DNA methylation levels at the cell cycle community CpGs in pancreatic cancer in TCGA (n=177). Histopathological features such as tumor stage at diagnosis, MKI67 expression (a marker of proliferation), and gender are annotated in each column. (B-C) Box plots showing the average DNA methylation levels at the cell cycle community CpGs in pancreatic cancer tumors from the TCGA cohort and PDAC data obtained from an independent cohort of PDAC tumors (Nones et al. [51]) respectively. Wilcoxon test p-values are denoted. DNA methylation levels in the PANC-1 pancreatic cancer cell line are also included. Bar plots showing the enrichment of the cell cycle community CpGs in (C) Segway-defined regulatory regions and (D) UniBind-defined TFBRs. The X-axis shows the log-transformed p-values obtained by hypergeometric testing using the 450k CpGs as background. (E) Hierarchical clustering of the cell cycle community gene expression levels in pancreatic cancer samples from TCGA (n=177). Histopathological features such as tumor stage at diagnosis, MKI67 expression, and gender are annotated. (F-G) Box plots showing the expression levels of the cell cycle community genes in pancreatic cancer in tumors from TCGA and Sandhu et al. respectively [54]. Wilcoxon test p-values are denoted. (H) Bar plot showing the enrichment of the emQTL in IM-PET-defined and GeneHancer-defined loops by emQTL community. Statistically significant enrichments are denoted with an asterisk. (I) Scatterplot showing the correlation between DNA methylation at the cell cycle bicluster CpGs and the expression of the cell cycle community genes in IM-PET loops in pancreatic cancer (TCGA-PANCAN, n=177). The Pearson correlation coefficient and the p-value are denoted.

#### The metabolism community

Among the cancer types with a correlation between DNA methylation at CpGs and expression of genes in the metabolism community was liver cancer. Moreover, this cancer type exhibited the lowest DNA methylation at the CpGs and expression of the genes in the metabolism community. Moreover, the CpGs in this community were significantly less methylated at the CpGs, and the genes were significantly expressed in this cancer type compared to normal tissue for this community. Therefore, we sought to investigate the role of the pan-cancer emQTLs in the metabolism community in further detail in this cancer type. By performing unsupervised hierarchical clustering of the DNA methylation levels of the CpGs and expression levels of the genes in the metabolism community, we observed clear heterogeneity in the expression profiles and DNA methylation profiles in liver cancer (Supplementary figure 16A and B respectively). According to the methylation data from TCGA, LICH tumors were significantly less methylated compared to normal adjacent tissue (Wilcoxon p-value=1.45e-10). They resembled methylation levels close to the purified HepG2 liver cancer cell line (Supplementary figure 16C). Similarly, the expression levels differed between liver cancer and normal adjacent tissue in which liver cancer had significantly lower expression levels of the genes in the metabolism community (Supplementary figure 16D, Wilcoxon p-value=2.55e-24). To assess in a cancer-specific manner whether the CpGs in the metabolism community were found in active chromatin, we obtained genome annotation data from Segway from the HepG2 cell line [55]. We found the CpGs in the metabolism community to be significantly enriched in regions carrying active enhancer and promoter chromatin marks. Next, we sought to investigate whether the emQTL in the metabolism community was enriched in predicted chromatin loops specific for a liver cancer cell line, and we found the emQTL in the metabolism community to be significantly enriched in IM-PET interaction loops (p-value=2.75e-45; Supp. table 1F). Altogether, our findings suggest that DNA methylation at enhancer and promoter regions regulating genes within the metabolism community is an important regulator of metabolism in several cancer types including liver cancer.

#### The hormone-like community

The cancer-type with the strongest negative median correlation coefficient between DNA methylation at the CpGs and expression of genes in the hormone-like community was pancreatic cancer. We continued to study the hormone-like community of this cancer type in further detail. Having found that the CpGs in the hormone-like community were enriched in the TFBR of TFs related to hormone-signaling, we further performed unsupervised hierarchical clustering of the DNA methylation levels at the CpGs and expression of the genes in the hormone-like community in pancreatic cancer. The DNA methylation profiles and expression profiles differed between the pancreatic tumor samples (Supplementary figure 17A-B). We found the CpGs in the hormone-like community to be significantly less methylated in tumor samples compared to normal adjacent tissue in two independent cohorts (Supplementary figure 17C-D; p-value=0.0278 and 1.27e-09 in TCGA and expression data obtained from Nones et al. [51] respectively). Moreover, the genes were significantly upregulated in TCGA and in a pancreatic cancer dataset obtained from Sandhu et al. [54] (p-value=0.00984 and 3.62e-05 respectively; Supplementary figure 17E-F). To assess the functional role of the CpGs in pancreatic cancer specifically with regard to histone modifications and regulatory regions, we performed an enrichment analysis using Segway-defined genome annotation data obtained from the PANC-1 [52, 53]. Our analysis showed significant enrichment of the hormone-related CpGs in enhancer and promoter regions in pancreatic cancer (Supplementary figure 17G; p-value=1.44e-165 and 3.29e-25 respectively; Supp. table 1H). Significant enrichment in IM-PET loops for the pancreatic cancer cell line PANC-1 was observed (Supplementary figure 17H; p-value=1.61e-65; Supp. table 1F). Taken together, these results suggest that the depletion of enhancer methylation at CpG sites in the hormone-like community may have a role in regulating transcriptional networks linked to hormone-signaling in several cancer types including pancreatic cancer.

## Discussion

Here, we performed for the first time emQTL analysis in a pan-cancer manner to identify disease-driving transcriptional networks dysregulated due to genome-wide loss of DNA methylation in cancer. We identify robust associations between DNA methylation levels at CpGs and gene expression that may contribute to alterations in transcriptional activity and cancer physiopathology; besides, some associations between DNA methylation and expression characterize the heterogeneity of the tumor microenvironment.

Using bipartite network analysis to group the pan-cancer emQTL, we identified six communities. Each community is composed of CpGs and genes for which methylation and expression levels are most often correlated. We showed that the six communities have a prognostic value for cancer patients’ survival. The communities highlight specific biological signals, and downstream analyses determined that the communities represent either heterogeneity of the cancer epigenome and transcriptome or varying levels of non-cancer cell infiltration. Three communities were related to alterations in cancer cells: Community 1 was linked to metabolism, Community 2 was linked to the regulation of proliferation, and Community 4 was linked to hormone-signaling.

For the cell cycle community, through the integration of ATAC-seq, ChIP-seq, and GeneHancer loop data, we show that loss of DNA methylation at enhancer regions at the TFBR of proliferation-related TFs is linked with a concomitant increase in expression of proliferation-related genes through GeneHancer-inferred interactions in most cancer types. Among the proliferation-related TFs were FOSL1/2 and JUN which can form AP-1 dimers previously shown to directly control the expression of cell cycle regulators such as cyclin D1, p53, p21, ARF, and p16 [56]. Among the cancer types, pancreatic cancers displayed the strongest emQTL correlations between DNA methylation at the CpGs and gene expression of the emQTL in the cell cycle community. Previous studies have shown that aberrant DNA methylation in pancreatic cancer is associated with aggressive phenotypes [51, 57–59]. Moreover, aberrant DNA methylation has been altered in genes related to core signaling pathways such as cell cycle regulation, Wnt/Notch signaling, and cell adhesion in pancreatic cancer [51, 60, 61]. Less is known about how the loss of methylation at distal regulatory regions influences pancreatic cancer tumor phenotype and prognosis. Our findings suggest that loss of DNA methylation is an important contributor to the upregulation of proliferation-related genes in pancreatic cancer. Also, our findings suggest that this finding applies to most cancer types.

The second cancer-related emQTL community was found to be related to hormone signaling in cancer. We previously described a hormone-like community related to estrogen signaling [19]. This study highlighted that DNA methylation plays a key role in regulating estrogen signaling in breast cancer. Our results suggest that DNA methylation might also play a key role in regulating hormone-signaling in other cancer types. We observe a significant loss of DNA methylation at the hormone-like community CpGs with a concomitant upregulation of genes in the hormone-like community in several cancer types. In pancreatic cancer, we observe that the CpGs are found in regions with active chromatin marks of enhancer regions. These enhancers are linked to the hormone-like genes through looping. Moreover, according to the TF enrichment analysis, the CpGs enriched in the binding site of TFs involved in hormone-signaling such as ESR1 [36] and AR [37]. Activation of AR associated with human carcinogenesis in pancreatic cancer has been well described [62]. Expression of estrogen receptor in pancreatic adenocarcinoma has previously been detected and has been linked to poor prognosis [63, 64]. Our findings suggest that loss of enhancer methylation also can be a driver of hormone-signaling in several cancer types such as pancreatic cancer.

The third cancer-related emQTL community was related to the regulation regulating metabolism in cancer. Reprogramming of the cellular metabolism is considered one of the core hallmarks of cancer, but little is still known about the mechanisms that drive this shift in the metabolism [1]. The CpGs were enriched for enhancers and binding regions of TFs involved in metabolism such as the HNF family of TFs [26–28] and RXRA [29]. Moreover, we found the emQTL to be more than expected by chance in GeneHancer and IM-PET loops. We thus hypothesize that loss of DNA methylation at promoter and enhancer regions may play a regulatory role on metabolism in cancer. More studies on the metabolism emQTL community will be of future interest.

In this study, we identify cancer-related emQTL representing associations between DNA methylation at CpGs and gene expression that may have varying effects on their cancer-promoting abilities. Some emQTL may be found in the periphery of regulatory regions and TFBSs and mimic the surrounding regions’ methylation levels. In contrast, others may be centrally located within the binding motifs of TFs and may therefore have a greater impact on gene transcription by blocking TF binding directly. Using the knowledge-guided emQTL approach, we identified candidate emQTL-CpGs that could have the greatest impact on the regulation of metabolism, proliferation, and hormone-signaling. We found that many candidate emQTL showed correlations within several cancer types. Our findings suggest that these candidate emQTL represent promising targets for targeted epigenetic treatment in the future that might be applied across cancer types.

We discovered three emQTL communities representing heterogeneity in the tumor cell type of the tumor microenvironment. One of these communities was the immune community which was found to be linked with varying infiltration of immune cells in the tumor tissues. The DNA methylation levels at CpGs of the immune community were negatively linked to immune infiltration; the methylation profiles of the tumors with high immune cell infiltration were closely related to the profiles of purified immune cells including T-cells, B-cells, leukocytes, and monocytes. Using tumor purity estimates based on copy number data, we could demonstrate the most likely cause of the immune community. This study emphasizes the significance of a better understanding of how cell types of the tumor microenvironment in bulk tissue influence gene expression and DNA methylation profiles obtained from tumors and signifies the importance of considering this concerning biomarker discovery.

Similarly, to the immune community, we found the EMT community to be linked with heterogeneity in the tumor microenvironment. In a previous study, we found varying infiltration of fibroblasts to be associated with EMT-related emQTL in breast cancer [18]. Fibroblasts are prominent cell types of the tumor microenvironment in solid tumors and express the genes found in the hallmark EMT gene set and may induce EMT in epithelial cells [65, 66]. Interestingly, among the genes residing in the EMT community were established fibroblast marker genes including COL1A2, COL5A1, and CD248 [67, 68]. We found a positive association between the xCell fibroblast score and the expression of the EMT community genes. Moreover, a negative association was found between DNA methylation and the xCell-derived fibroblast score. The tumors with the highest fibroblast infiltration had methylation profiles resembling the methylation values obtained from purified fibroblasts. Fibroblast infiltration was also negatively associated with tumor purity. Altogether our findings suggest that while the EMT community carries EMT features, one of the most probable causes of this community is the presence of fibroblasts in the tumors. Fibroblast infiltration has previously been linked to poor treatment response in several cancer types and has been found to modulate immune cell activity and promote immune evasion [69]. Moreover, an increasing number of studies have found fibroblast infiltration to promote tumorigenesis through the secretion of cancer-associated fibroblast (CAF) secreted factors such as TGF-β [70]. TGF-β stimulation has been shown to induce changes in the DNA methylation landscape in several cancer types such as ovarian cancer, liver cancer, and prostate cancer [71–73]. Interestingly, the TGFB gene was found residing within the EMT community, suggesting a possible role in cancer carcinogenesis either directly by upregulation in the cancer cells or indirectly through crosstalk between CAFs and tumor cells. Although the EMT community is associated with fibroblast infiltration, we further hypothesize that there could be a less detectable EMT signal in the EMT community from the tumors themselves that is masked by fibroblast infiltration in the tumor biopsies. As studies have shown, crosstalk between fibroblasts and cancer cells through CAF-secreted factors which can alter the methylation landscape of cancer cells and promote EMT [70–73]. A more detailed study on the EMT community regarding EMT in cancer cells would therefore be of future interest.

In contrast to the immune and EMT communities, we found the NE-like community to be partly explained by differences in the tissue of origin and heterogeneity in the TME. The NL tumors had distinct methylation and expression profiles compared to the other TCGA cancer types. The distinct methylation and expression profiles concerning the NE-like community in the NL and NNL tumors suggest that the cell of origin is the major driver of the negative correlation in the NE-like community. However, since the methylation levels at the CpGs and expression of the genes also deviated between the normal and tumor samples in NNL, the cell of origin only partly explains this correlation. We assessed the link between the DNA methylation at CpGs and the expression of the genes in the NE-like community in NL and NNL tumors independently. We found a significant association with the tumor microenvironment score derived from xCell, which is a combined score of the stromal and immune scores. In summary, our findings suggest that the NE-like community is caused by differences in the tissue of origin between the tumors and that differences in tumor composition cause variability in gene expression and DNA methylation within the cancer-types.

## Conclusion

Through our pan-cancer emQTL approach, we provide a comprehensive view of the epigenetically-related mechanisms in cancer. We identify novel emQTL networks related to cancer-associated transcriptional networks including proliferation, metabolism, hormone-signaling, and infiltration of cell types of TME such as fibroblasts and immune cells. Furthermore, genome-wide loss of enhancer methylation at specific CpG sites was linked with transcriptional dysregulation of genes related to proliferation, metabolism, and hormone-signaling in cancer.

Our study provides insight into how alterations in DNA methylation in the context of histone modifications and the chromatin 3D structure govern and influence transcriptional regulation in a cancer-specific manner. Our knowledge-guided emQTL approach identifies potential therapeutic targets for future targeted epigenetic treatment and cancer biomarkers. Furthermore, we highlight the impact of non-tumor cell infiltration on epigenetic and transcriptomic profiles in tumors.

## Material and methods

### Patient material and data processing

The Cancer Genome Atlas Program (TCGA) has previously been described [74]. Level 3 DNA methylation (beta values), ATAC-seq (log2((count+5)PM)-qn), and gene expression (log2(fpkm-uq+1)) data from the TCGA-PANCAN dataset were downloaded from the UCSC Xena browser [75]. Probes with more than 50% missing values were removed and the remaining missing values were imputed using the R package impute v1.70.0 (function *impute.knn*) for each cancer type independently. For each cancer type, the number of neighbors used in the imputation (k) was set to 10 except for ovarian cancer in which k was set to 3 due to few samples with DNA methylation data from the Illumina HumanMethylation 450k array available.

DNA methylation and gene expression data obtained from primary pancreatic cancer tumors and normal tissue used for validation of the findings in cell cycle- and hormone-like community was obtained from Nones et al. (GEO accession number GSE49149; Illumine Human Methylation 450k) [51] and from Sandhu et al. (SurePrint G3 Human GE 8×60k microarrays)[54] respectively.

### Cell line data

Raw intensity (IDAT) files with methylation data from the HepG2 and PANC-1 cell line were retrieved from encode with accession codes GSE128685 and GSE40699 respectively. Beta-values were imputed using the minfi R package v1.42.0 (function *getBeta*). DNA methylation data with pre-calculated beta values were obtained from: T-cells (GSE79144), monocytes (GSE68456), leukocytes (GSE69270), B-cells (GSE68456), MDAMB231 (GSE94943) and MCF7 (GSE69188). DNA methylation data from the NCI-60 cell lines were downloaded from the CellMiner database (https://discover.nci.nih.gov/cellminer/home.do) [76].

### Statistical and bioinformatic analyses

All data analyses were performed using the R software version 4.0.2 [77] unless otherwise specified. A p-value<0.05 was considered statistically significant in this study. The interaction plot between the emQTL communities was generated using Cytoscape v3.9.1 in which all 192 links from community 4 and 4808 randomly selected emQTL from communities 1-5 were included. Random links were selected by setting the seed parameter to 42 using the R function *set.seed*. Each community was labeled using the AutoAnnotate v1.4.0 app in Cytoscape. Pan-cancer emQTL analysis R code is available on GitHub (https://github.com/JorgenAnkill/Pan-cancer-expression-methylation-Quantitative-Trait-Loci-analysis). UpSet plots were made using the UpSet R package v1.4.0 [78].

### Genome-wide correlation analysis

The TCGA pan-cancer tumor samples with matching DNA methylation and gene expression data available were randomly split into a discovery and validation dataset (R function *sample*, seed=42). All CpGs with an interquartile range (IQR)>0.1 (191 636 CpGs) and genes with a non-zero IQR (17 265 genes) in the discovery dataset (*n*=4104) were tested for significant correlations using Pearson’s correlation. Due to the large number of associations, we only included those with an absolute Pearson correlation coefficient>0.5 (p-value<1.40e-258) and a Bonferroni-corrected p-value less than 0.05 (p-value<1.51e-11). for downstream analyses. Significant CpG-gene associations from the discovery dataset were reanalyzed in the validation dataset (*n*=4089). Associations were validated if the absolute correlation coefficient<0.5 and had a Bonferroni-corrected p-value less than 0.05 (p-value<3.67e-08). In this study, we only focused on the negative correlations in subsequent analyses.

### Global pan-cancer emQTL network community detection using CONDOR

A clustering algorithm for community detection was applied to the pan-cancer emQTL with negative correlations using the *condor* R package v1.1.1 (function *condor.cluster*) [20]. The multilevel community was set as the clustering method and the absolute Pearson correlation coefficient of the pan-cancer emQTL was used as weights. For reproducibility, the initial state of the generator (seed) was set to 42.

### Gene set enrichment analysis

Gene sets used for GSEA were obtained from the Molecular Signatures Database (MSigDB) v.7.4 [21]. Enrichment of genes in each gene set of interest was performed by hypergeometric testing using the C5 gene ontology- and hallmark (H) gene set collections. Enrichment with a BH-corrected p-value<0.05 was considered statistically significant.

### Hierarchical clustering of DNA methylation and gene expression levels

Clustering of DNA methylation and expression data was performed using the R package *pheatmap* v1.0.12 with Euclidean distance and ward.D2 clustering method unless otherwise specified. Expression values were transformed into z-scores before clustering for visualization purposes.

### Genomic annotation

Genome annotation data of *cis*-Regulatory Elements (cCREs) from The Encyclopedia of DNA Elements (ENCODE) [24] was downloaded from the SCREEN webtool (https://screen.encodeproject.org). The registry encompasses 1,063,878 human regulatory regions from 4834 experimental datasets representing 1,518 different cell types and tissues. Nine different cCREs were included from the dataset; “pELS, CTCF-bound”, “dELS, CTCF-bound”, “PLS, CTCF-bound”, “CTCF-only, CTCF-bound”, ”dELS”, ”DNase-H3K4me3, CTCF-bound”, “pELS”, “PLS” and “DNase-H3K4me3”. In downstream analyses, some of these annotations were collapsed/converted into one as follows: Proximal CTCF-bound enhancer=”pELS, CTCF bound”, Distal CTCF-bound enhancer=” dELS, CTCF-bound”, CTCF-bound promoter=” PLS, CTCF-bound” and “DNase-H3K4me3, CTCF-bound”, Promoter =” PLS” and “DNase-H3K4me3”, Distal enhancer=” dELS”, pELS=” Proximal enhancer” and CTCF=” CTCF-only, CTCF-bound”.

Cell-type specific genome-wide segmentation data from pancreatic cancer cell line PANC1 and the liver cancer cell line HepG2 was retrieved from Segway (https://segway.hoffmanlab.org). Segway is a method used to annotate the genome using a dynamic Bayesian network model on ChIP-seq data obtained from cell lines and/or tissues [53]. Enrichment of emQTL-CpGs in Segway-defined regulatory regions was assessed by hypergeometric testing (R function *phyper*) using the IlluminaMethylation450k CpGs as background. Adjusted p-values were obtained by Benjamini-Hochberg correction.

### Transcription factor enrichment analysis

Maps of direct TF-DNA interactions were obtained from the UniBind database (https://unibind2018.uio.no) [25]. The UniBind TFBSs correspond to high-confidence direct TF-DNA interactions determined experimentally through ChIP-seq and computationally through position weight matrices (PWMs) from JASPAR [79]. The predicted TFBSs were derived from 1983 ChIP-seq experiments for 231 TFs across cell types and tissues and were predicted to have high PWM scores and be near ChIP-seq peaks.

Since the genomic coordinates of the UniBind database are based on the hg38 reference genome, all CpG positions from the Illumina 450k array were lifted over from hg19 to hg38 using the LiftOver web tool from the UCSC genome browser (https://genome.ucsc.edu) and were extended ±150bp from TFBSs. The CpG positions were then intersected with the TBFR using BEDTools v2.27.1 [80]. Since UniBind maps of TF binding sites are often derived from several ChIP-seq experiments for each TF, we merged the TF binding sites for all ChIP-seq experiments for each TF. Enrichment of CpGs in TFBRs was imputed using hypergeometric testing (R function *phyper*) using the IlluminaMethylation450k CpGs as background. Multiple testing was accounted for using Benjamini-Hochberg correction (R function *p.adjust*).

### Cell type enrichment analysis by xCell

Deconvolution of the cellular composition of the tumor samples in the TCGA-PANCAN dataset was performed using the xCell [49] r-package version 1.1.0 (function *xCellAnalysis*). xCell is a machine learning algorithm trained on 64 stromal and immune datasets to identify specific cell types from bulk tissue using 10 808 genes as signatures. To assess the link between the xCell-derived cell type scores, the tumor samples were divided into quartile groups based on the xCell score; i.e., the level of infiltration. Differences in DNA methylation at CpGs or expression of genes between the groups were then assessed using the Kruskal-Wallis test.

### Predicted chromatin interactions

Cis-regulatory elements-gene associations were obtained from the GeneHancer database v5.0 [44]. Enhancer to gene associations was obtained from GeneCards Suite Version 5.0 [44]. The genomic coordinates were lifted over from hg38 to hg19 using the LiftOver web tool from the UCSC genome browser (https://genome.ucsc.edu). Predicted Integrated Methods for Predicting Enhancer Targets (IM-PET) interactions from the PANC-1, HepG2, A549, HCC1954, MCF7, HCT116, HELA, and K562 cell lines were retrieved from the 4D Genome data portal (https://4dgenome.research.chop.edu) [81]. IM-PET is a computational algorithm used to predict chromatin interaction loops between enhancers and promotors by using genomic features including distance restrictions, enhancer-target promoter activity profile correlation, TF and target promoter correlation, and co-evolution of enhancer and target promoter [82]. BEDTools v2.27.1 was used to intersect gene and CpG positions with the genomic intervals defining the feet of the chromatin loops for the IM-PET and GeneHancer data. An emQTL was considered in a chromatin loop if the CpG and gene were found in different feet of the same chromatin loop. Enrichment of emQTL in chromatin loops was calculated using hypergeometric test (R function *phyper*) using all possible *in cis* (i.e., on the same chromosome) pairs between genes and CpGs in the 450k array as background. Visualization of the chromatin interactions was made using the *Gviz* v1.40.1 [83] and *GenomicRanges* v1.48.0 [84] R packages. Genomic interactions were visualized in Gviz using the *GenomicInteractions* v1.30.0 [85] R packages.

### Tumor purity data

Allele-Specific Copy Number Analysis of Tumors (ASCAT) [48] is a computational tool that estimates tumor purities by inferring the fraction of tumor cells in a sample. This is done by analyzing the proportion of tumor-specific and normal-specific allele frequencies in DNA copy number data. ASCAT compares the observed DNA copy number data to a reference profile, taking into account DNA contamination, ploidy of cancer cells and the presence of subclonal populations of cells with different copy number alterations. Tumor purity estimates for the TCGA samples were obtained from the NCI Genomic Data Commons (GDC) data portal upon request (https://portal.gdc.cancer.gov) [86].

### Assessing the predictive power of the pan-cancer emQTL

The ability of the pan-cancer emQTL to predict prognosis was performed by building Cox regression with ridge penalty to contract models for survival predictions by estimating the hazard function (Ridge Cox model) [45]. Specifically, PCA was performed on the z-scores obtained from methylation and expression data for the emQTL-CpGs and genes. In each community, the first two principal components (PCs) of emQTL-CpGs and the first two PCs of the emQTL-genes were extracted; 24 PCs were extracted from the six communities. We regressed patients’ survival in each cancer type on the 24 PCs using the Ridge Cox model. The survival prediction metric Uno’s C-index was obtained using repeated 5-fold cross-validation to overcome overfitting.

Kaplan-Meier survival curves and log-rank tests were performed using the R function *Surv* and *survfit* from the R package *survival* v3.5.3. Principal component analysis (PCA) was performed on the z-scores of the expression and methylation data for each cancer type in TCGA using the R package *stats* v3.6.2. The principal components were then used as covariates in Ridge Cox models for each cancer type using the R package *glmnet* v4.1.6. Uno’s C-index was then estimated using the R package *survAUC* v1.1.1 and summarized based on repeated 5-fold cross-validation. The seed parameter was initialized as 123 using the R function *set.seed*. Survival plots were made using the *survminer* R package v0.4.8.

### Knowledge-guided emQTL analysis

Knowledge-guided emQTL was performed based on several criteria related to the biological function of the cancer-related emQTL communities (i.e., not caused by infiltration). Since all cancer-related community CpGs were most significantly enriched in enhancer regions, the CpG-gene pair of the cancer-promoting candidate emQTL had to be (I) on opposite sides of predicted chromatin interaction loops according to GeneHancer, and (II) be located in one of the significantly enriched enhancer segments in the ENCODE Portal database. (III) The CpG must be in the binding region of one of the enriched TFs for the community with the corresponding biological process of interest (i.e., metabolism, proliferation, or hormone-signaling) as defined by UniBind, and (IV) the gene must be a part of a curated gene set associated with the biological process of interest. The DNA methylation levels at the CpGs and expression of genes fulfilling these criteria were then correlated in the discovery dataset before validating the association in the validation dataset. Only candidates with a Bonferroni corrected p-value<0.05 that could be validated in the validation dataset were considered candidate emQTL.

## Data material and availability

R code related to the pan-cancer emQTL analysis is available at GitHub (https://github.com/JorgenAnkill/Pan-cancer-expression-methylation-Quantitative-Trait-Loci-analysis). DNA methylation, ATAC-seq and gene expression data from the PAN-CAN dataset can be obtained from the UCSC Xena browser data portal (https://xenabrowser.net) [75]. DNA methylation data from pancreatic ductal adenocarcinomas were obtained from GEO with accession number GSE49149 [51]. Maps of TF-DNA interactions used in this study can be downloaded from the UniBind database (https://unibind2018.uio.no). DNA methylation data from several cell lines are available from GEO: T-cells (GSE79144), monocytes (GSE68456), leukocytes (GSE69270), B-cells (GSE68456), HepG2 (GSE128685), PANC-1 (GSE40699), MDAMB231 (GSE94943) and MCF7 (GSE69188). DNA methylation data from the NCI-60 cell lines were downloaded from the CellMiner database (https://discover.nci.nih.gov/cellminer/home.do) [76]. Genome-wide annotation data from Segway was obtained from (https://segway.hoffmanlab.org) [52] and breast cancer subtype-specific ChromHMM segmentation data was obtained from Xi et al. [87]. GeneHancer loops were obtained from GeneCards Suite Version 5.0 [44]. Genome annotation data of cCREs were downloaded from SCREEN (https://screen.encodeproject.org) [24]. IM-PET data from the HCC1954, PANC-1, A549, MCF7, HCT116, HELA, K562 and the HepG2 cell line was obtained from the 4D Genome data portal (https://4dgenome.research.chop.edu) [81]. Tumor purity estimates based on ASCAT were obtained from the GDC data portal (https://portal.gdc.cancer.gov).

## Supplementary table legends

**Supplementary Table 1. Summary of data from this study.** (A) Overview of the CpGs in the emQTL communities. (B) Overview of the genes in each emQTL community. (C) Enrichment of genes in gene sets from the MSigDB by community. Only gene sets with a significant enrichment after BH-correction are shown. (D) Enrichment of CpGs in community 1-6 in Encode-defined cis-regulatory elements defined by ENCODE. (E) Enrichment of CpGs in community 1-6 in UniBind-defined TFBRs. Only TFs showing a statistically significant enrichment are shown. (F) Enrichment of emQTL in community 1-6 in GeneHancer-defined chromatin loops and IM-PET loops. (G) Candidate cancer-promoting emQTL related to the regulation of metabolism, proliferation, and hormone-signaling. p-values and r-values obtained from Pearson correlation between DNA methylation at CpGs and expression levels of genes are included for the discovery and validation dataset from the TCGA-PANCAN dataset. R-values obtained from the correlation analysis in the 33 different TCGA-PANCAN cancer types independently are annotated if the correlation was significant after BH correction. (H) Enrichment emQTL-CpGs in Segway-defined regulatory regions in the PANC-1 and HepG2 cell lines by community.

## Acknowledgments

This work was supported by a grant from South-Eastern Norway Regional Health Authority (grant 2020031 to TF). I would also like to thank Elin Hegland Kure for kindly sharing expression data from pancreatic cancer from Sandhu et al. [54].

## Abbreviations

AR: Androgen receptor
ASCAT: Allele-Specific Copy Number Analysis of Tumors
BH: Benjamini-Hochberg
BRCA: Breast invasive carcinoma
ChIA-PET: Chromatin Interaction Analysis with Paired-End Tag
dELS: TSS-distal enhancer-like signatures
DNMT: DNA methyltransferase
emQTL: Expression-methylation Quantitative Trait Loci Analysis
ENCODE: The Encyclopedia of DNA Elements
EMT: epithelial-mesenchymal transition
ESR1: Estrogen receptor alpha
FE: Fold enrichment
FOXA1: Forkhead Box A1
GATA: GATA binding protein 2
GEO: Gene Expression Omnibus
GSEA: Gene set enrichment analysis
IM-PET: Integrated Methods for Predicting Enhancer Targets
NCI: National Cancer Institute
PAAD: Pancreatic adenocarcinoma
PCA: Principal component analysis
PC: Principal component
PDAC: Pancreatic ductal adenocarcinoma
pELS: TSS-proximal enhancer-like signatures
PLS: promoter-like signatures
TCGA: The Cancer Genome Atlas
TET: Ten-eleven translocation
TF: Transcription factor
TFBR: Transcription factor binding region
TFBS: Transcription factor binding site
TME: Tumor microenvironment

