## Supplementary figures for "Integrative pan-cancer analysis reveals a common architecture of dysregulated transcriptional networks characterized by loss of enhancer methylation"

This file contains:

- Supplementary Figure S1-S17

### SUPPLEMENTARY FIGURES

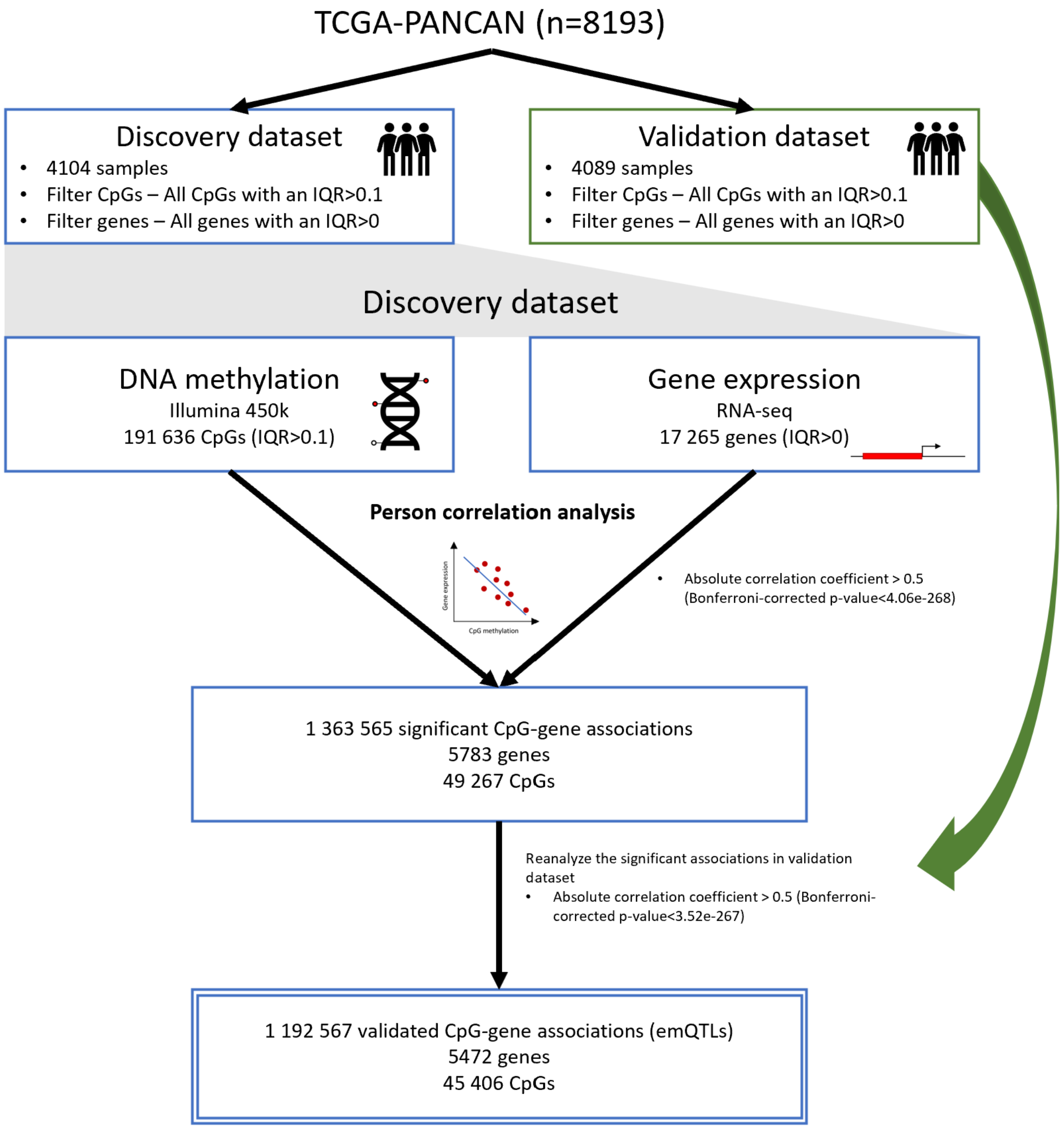


**Supplementary figure 1. Overview of the pan-cancer emQTL pipeline.** Assessment of the link between DNA methylation and gene expression was performed by Pearson correlation. The significant correlations found in the discovery subset of the TCGA-PANCAN dataset (n=4104) were then validated in the TCGA-PANCAN validation dataset (n=4089).

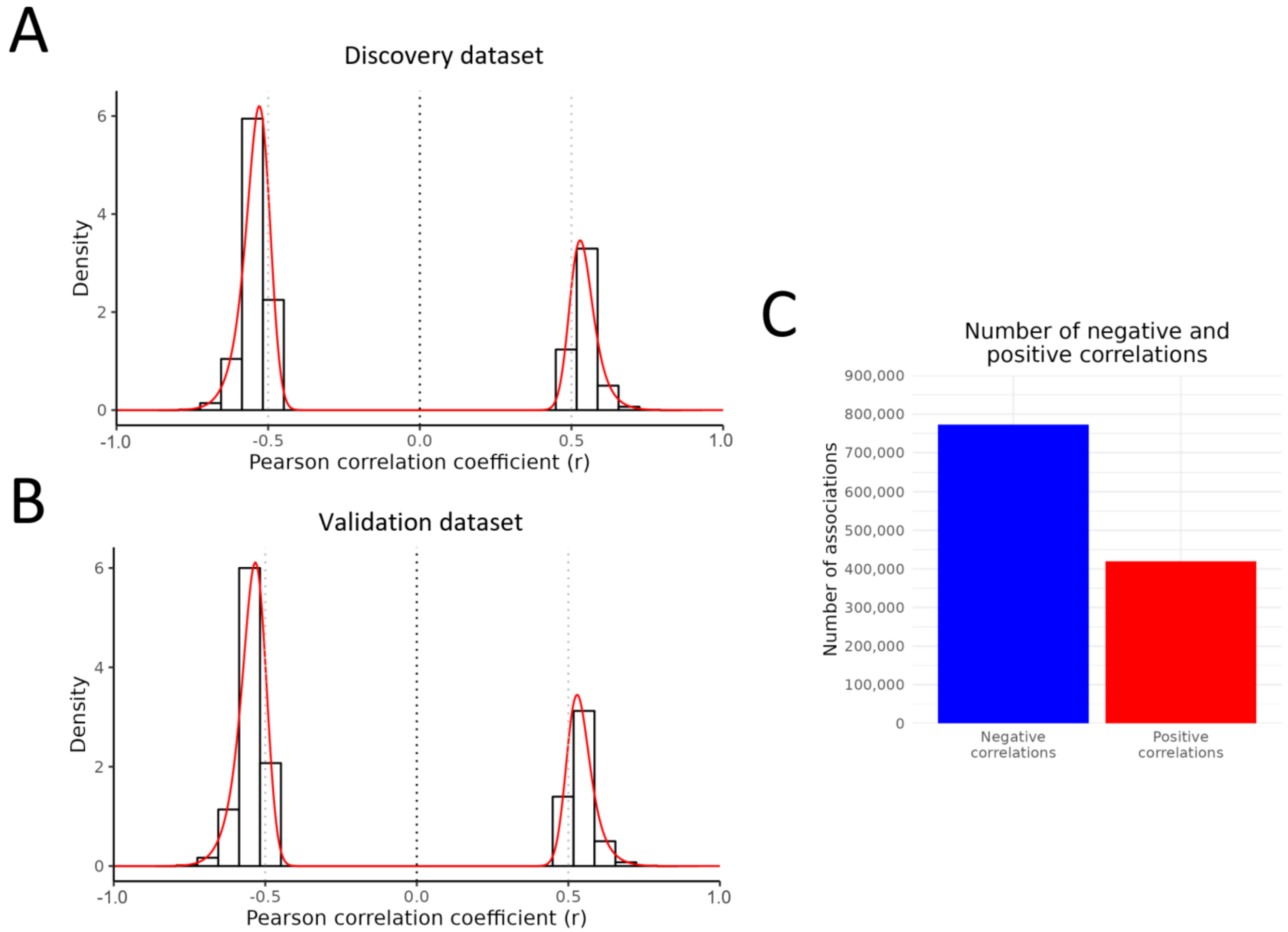


**Supplementary figure 2. Overview of the pan-cancer emQTL.** Density plot showing the distribution of the Pearson correlation coefficients in the discovery (A) and validation (B) datasets from the TCGA-PANCAN dataset for all the 1 192 567 pan-cancer emQTL. (C) Bar plot showing the number of pan-cancer emQTL showing negative and positive correlations between DNA methylation and gene expression.

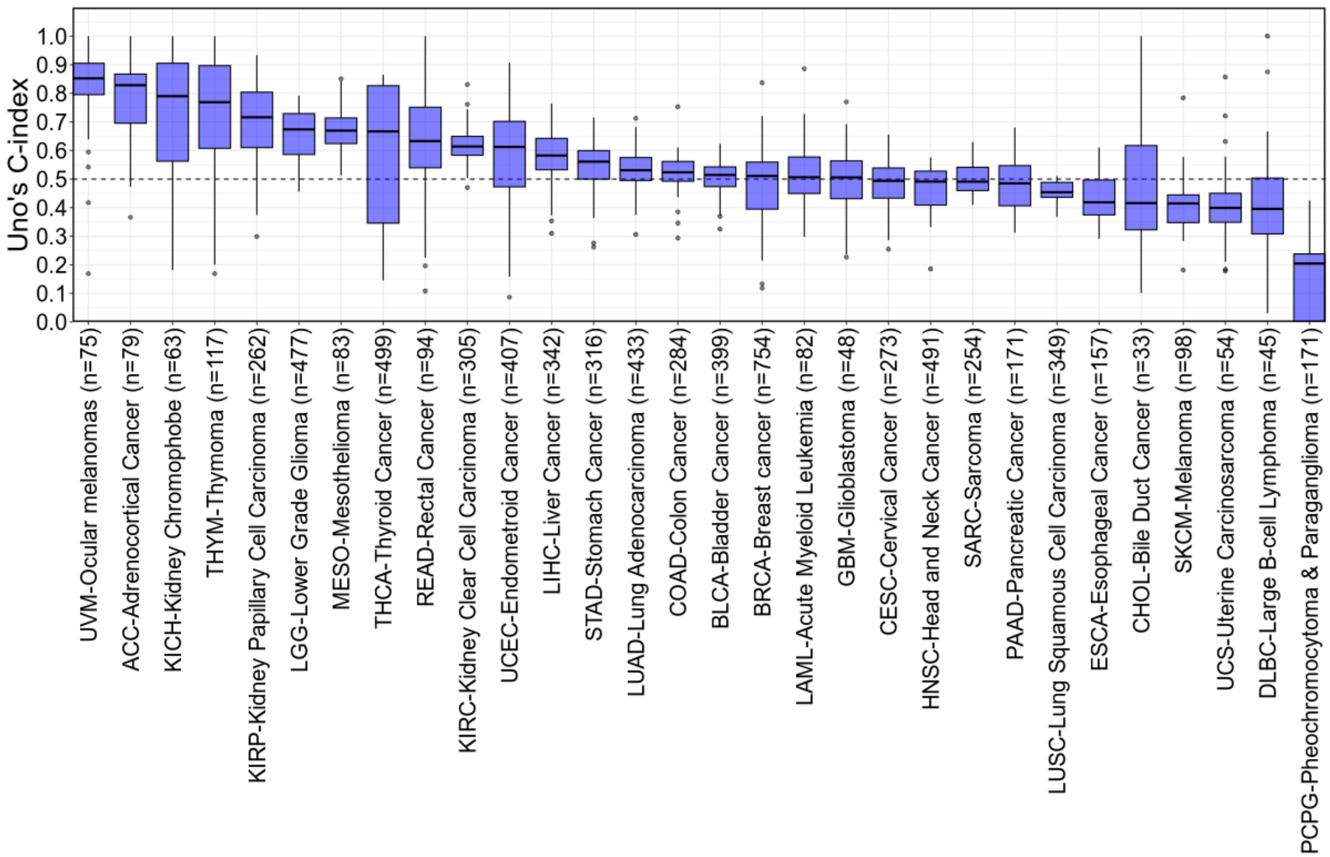


**Supplementary figure 3. Survival prediction of the pan-cancer emQTL.** (A) The pan-cancer patients were randomly split into 80% training and 20% test data 50 times. Each box represents Uno's C-indexes estimated from the randomly split 20% test data. Due to few samples for ovarian cancer (n=8) and high censoring in prostate- and testicular cancer, these cancer types were not included in the analysis.

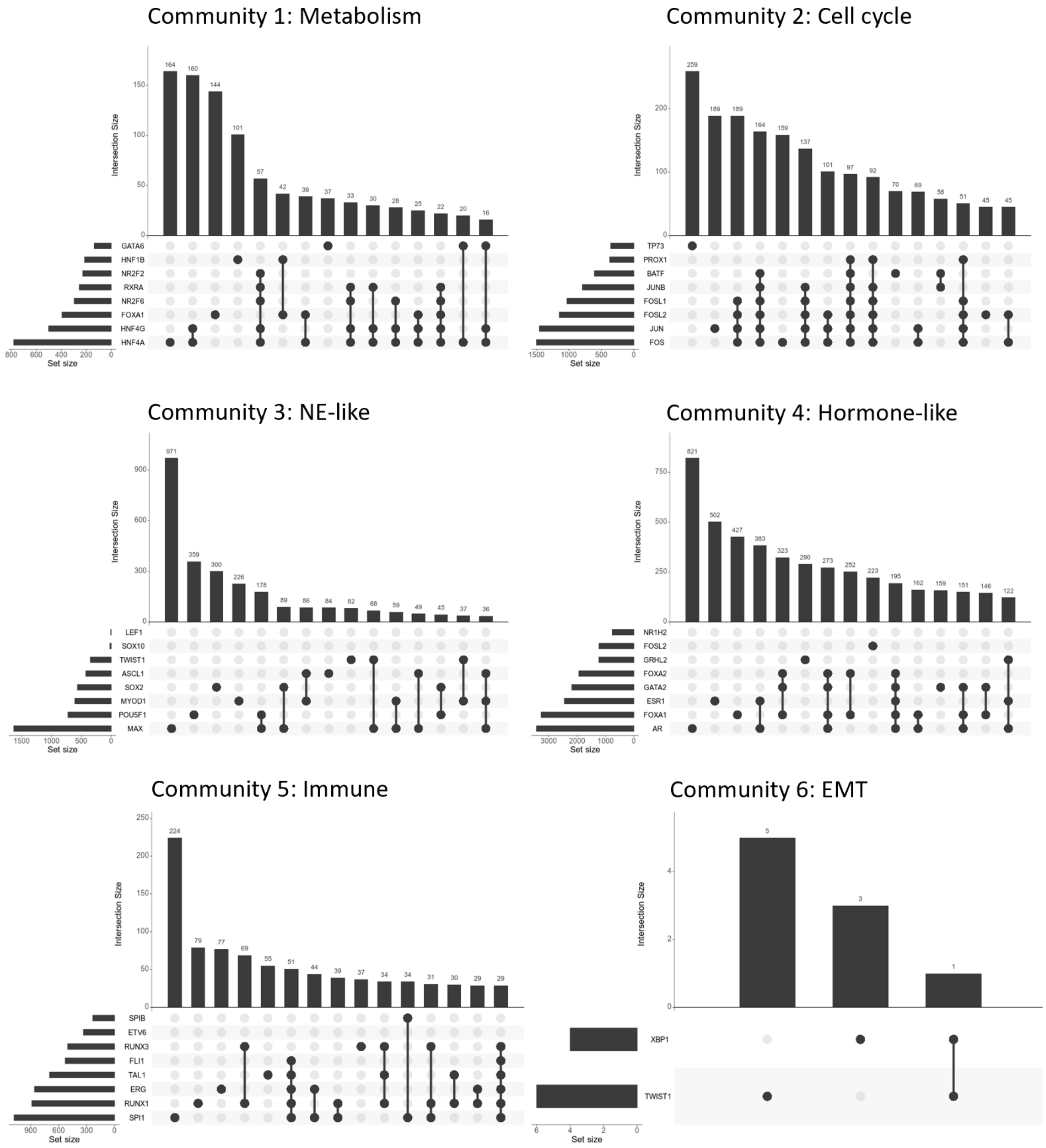


**Supplementary figure 4.** Overview of the TFBR overlap between the top enriched TFs in each emQTL community represented as Upset plots.

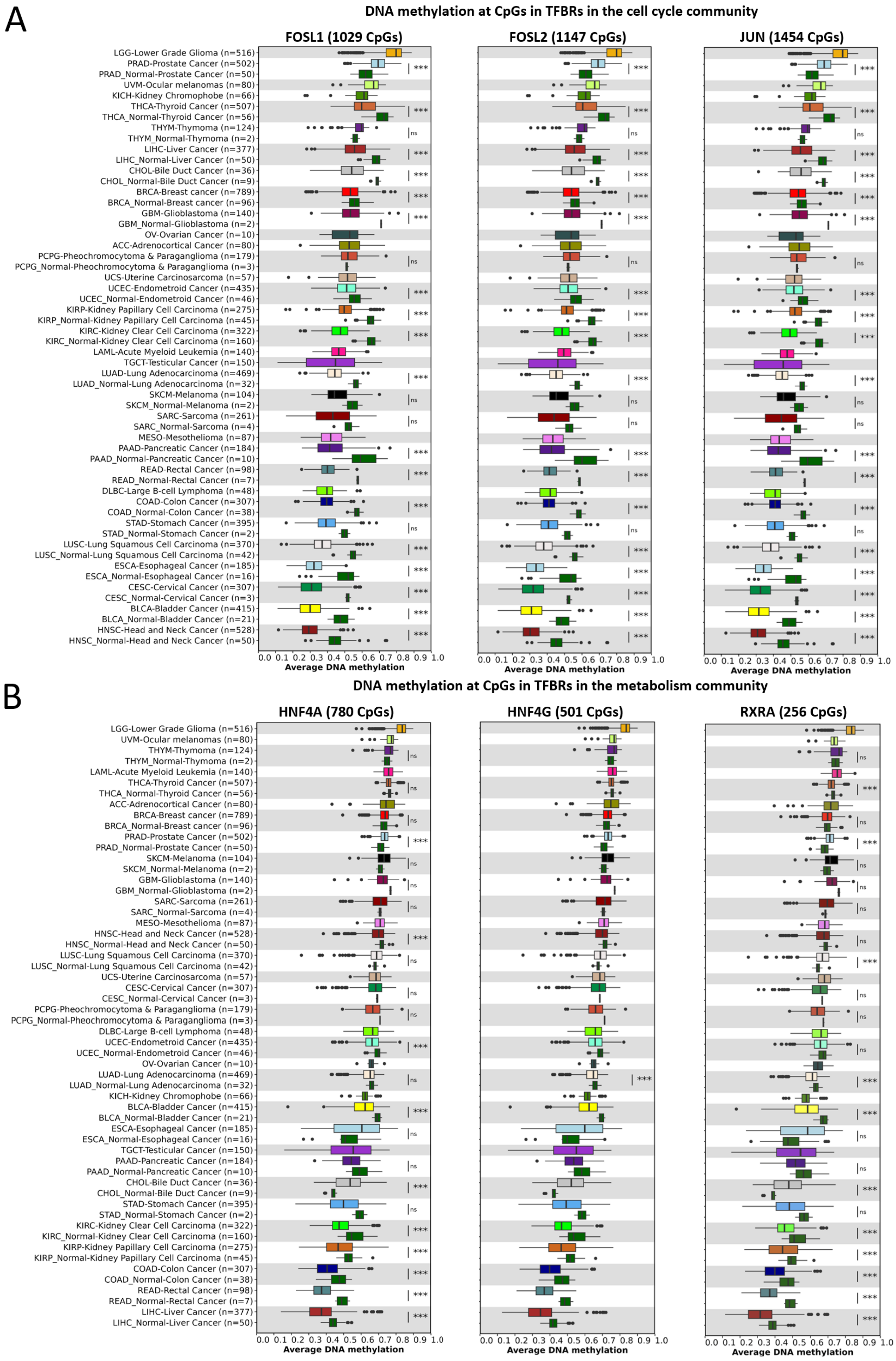
A
DNA methylation at CpGs in TFBRs in the cell cycle community
FOSL1 (1029 CpGs)
FOSL2 (1147 CpGs)
JUN (1454 CpGs)
LGG-Lower Grade Glioma (n=516)
PRAD-Prostate Cancer (n=502)
PRAD_Normal-Prostate Cancer (n=50)
UVM-Ocular melanomas (n=80)
KICH-Kidney Chromophobe (n=66)
THCA-Thyroid Cancer (n=507)
THCA_Normal-Thyroid Cancer (n=56)
THYM-Thymoma (n=124)
THYM_Normal-Thymoma (n=2)
LIHC-Liver Cancer (n=377)
LIHC_Normal-Liver Cancer (n=50)
CHOL-Bile Duct Cancer (n=36)
CHOL_Normal-Bile Duct Cancer (n=9)
BRCA-Breast cancer (n=789)
BRCA_Normal-Breast cancer (n=96)
GBM-Glioblastoma (n=140)
GBM_Normal-Glioblastoma (n=2)
OV-Ovarian Cancer (n=10)
ACC-Adrenocortical Cancer (n=80)
PCPG-Pheochromocytoma & Paraganglioma (n=179)
PCPG_Normal-Pheochromocytoma & Paraganglioma (n=3)
UCS-Uterine Carcinosarcoma (n=57)
UCEC-Endometroid Cancer (n=435)
UCEC_Normal-Endometroid Cancer (n=46)
KIRP-Kidney Papillary Cell Carcinoma (n=275)
KIRP_Normal-Kidney Papillary Cell Carcinoma (n=45)
KIRC-Kidney Clear Cell Carcinoma (n=322)
KIRC_Normal-Kidney Clear Cell Carcinoma (n=160)
LAML-Acute Myeloid Leukemia (n=140)
TGCT-Testicular Cancer (n=150)
LUAD-Lung Adenocarcinoma (n=469)
LUAD_Normal-Lung Adenocarcinoma (n=32)
SKCM-Melanoma (n=104)
SKCM_Normal-Melanoma (n=2)
SARC-Sarcoma (n=261)
SARC_Normal-Sarcoma (n=4)
MESO-Mesothelioma (n=87)
PAAD-Pancreatic Cancer (n=184)
PAAD_Normal-Pancreatic Cancer (n=10)
READ-Rectal Cancer (n=98)
READ_Normal-Rectal Cancer (n=7)
DLBC-Large B-cell Lymphoma (n=48)
COAD-Colon Cancer (n=307)
COAD_Normal-Colon Cancer (n=38)
STAD-Stomach Cancer (n=395)
STAD_Normal-Stomach Cancer (n=2)
LUSC-Lung Squamous Cell Carcinoma (n=370)
LUSC_Normal-Lung Squamous Cell Carcinoma (n=42)
ESCA-Esophageal Cancer (n=185)
ESCA_Normal-Esophageal Cancer (n=16)
CESC-Cervical Cancer (n=307)
CESC_Normal-Cervical Cancer (n=3)
BLCA-Bladder Cancer (n=415)
BLCA_Normal-Bladder Cancer (n=21)
HNSC-Head and Neck Cancer (n=528)
HNSC_Normal-Head and Neck Cancer (n=50)
0.0 0.1 0.2 0.3 0.4 0.5 0.6 0.7 0.8 0.9 1.0
Average DNA methylation
B
DNA methylation at CpGs in TFBRs in the metabolism community
HNF4A (780 CpGs)
HNF4G (501 CpGs)
RXRA (256 CpGs)
LGG-Lower Grade Glioma (n=516)
UVM-Ocular melanomas (n=80)
THYM-Thymoma (n=124)
THYM_Normal-Thymoma (n=2)
LAML-Acute Myeloid Leukemia (n=140)
THCA-Thyroid Cancer (n=507)
THCA_Normal-Thyroid Cancer (n=56)
ACC-Adrenocortical Cancer (n=80)
BRCA-Breast cancer (n=789)
BRCA_Normal-Breast cancer (n=96)
PRAD-Prostate Cancer (n=502)
PRAD_Normal-Prostate Cancer (n=50)
SKCM-Melanoma (n=104)
SKCM_Normal-Melanoma (n=2)
GBM-Glioblastoma (n=140)
GBM_Normal-Glioblastoma (n=2)
SARC-Sarcoma (n=261)
SARC_Normal-Sarcoma (n=4)
MESO-Mesothelioma (n=87)
HNSC-Head and Neck Cancer (n=528)
HNSC_Normal-Head and Neck Cancer (n=50)
LUSC-Lung Squamous Cell Carcinoma (n=370)
LUSC_Normal-Lung Squamous Cell Carcinoma (n=42)
UCS-Uterine Carcinosarcoma (n=57)
CESC-Cervical Cancer (n=307)
CESC_Normal-Cervical Cancer (n=3)
PCPG-Pheochromocytoma & Paraganglioma (n=179)
PCPG_Normal-Pheochromocytoma & Paraganglioma (n=3)
DLBC-Large B-cell Lymphoma (n=48)
UCEC-Endometroid Cancer (n=435)
UCEC_Normal-Endometroid Cancer (n=46)
OV-Ovarian Cancer (n=10)
LUAD-Lung Adenocarcinoma (n=469)
LUAD_Normal-Lung Adenocarcinoma (n=32)
KICH-Kidney Chromophobe (n=66)
BLCA-Bladder Cancer (n=415)
BLCA_Normal-Bladder Cancer (n=21)
ESCA-Esophageal Cancer (n=185)
ESCA_Normal-Esophageal Cancer (n=16)
TGCT-Testicular Cancer (n=150)
PAAD-Pancreatic Cancer (n=184)
PAAD_Normal-Pancreatic Cancer (n=10)
CHOL-Bile Duct Cancer (n=36)
CHOL_Normal-Bile Duct Cancer (n=9)
STAD-Stomach Cancer (n=395)
STAD_Normal-Stomach Cancer (n=2)
KIRC-Kidney Clear Cell Carcinoma (n=322)
KIRC_Normal-Kidney Clear Cell Carcinoma (n=160)
KIRP-Kidney Papillary Cell Carcinoma (n=275)
KIRP_Normal-Kidney Papillary Cell Carcinoma (n=45)
COAD-Colon Cancer (n=307)
COAD_Normal-Colon Cancer (n=38)
READ-Rectal Cancer (n=98)
READ_Normal-Rectal Cancer (n=7)
LIHC-Liver Cancer (n=377)
LIHC_Normal-Liver Cancer (n=50)
0.0 0.1 0.2 0.3 0.4 0.5 0.6 0.7 0.8 0.9 1.0
Average DNA methylation

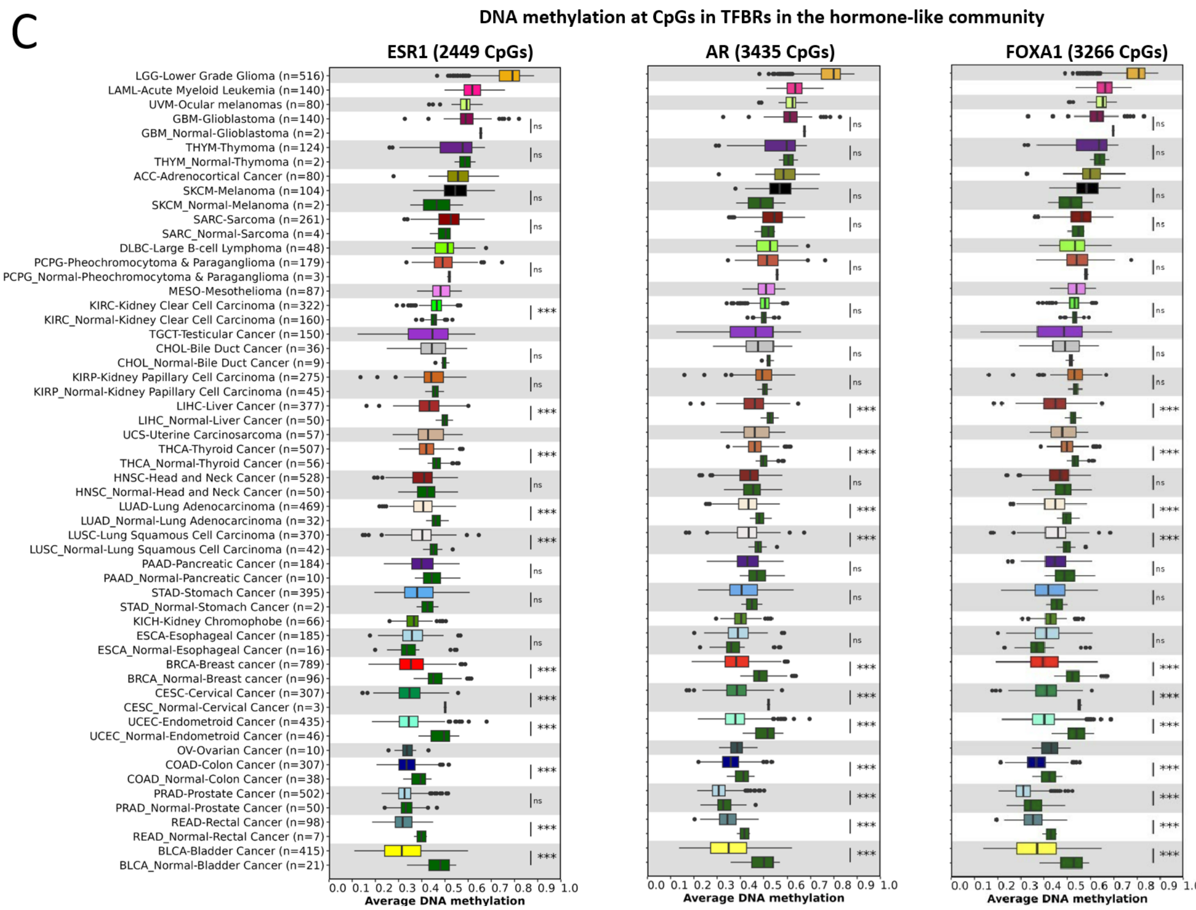


**Supplementary figure 5. DNA methylation at the TFBR in the cell cycle, metabolism, and hormone-like emQTL community.** Box plots showing the average DNA methylation at the CpGs in the TFBR of some of the top enriched TFs in the cell cycle (A), metabolism (B), and hormone-like (C) communities. DNA methylation levels from available normal samples are included. BH-corrected Wilcoxon-test p-values are denoted.

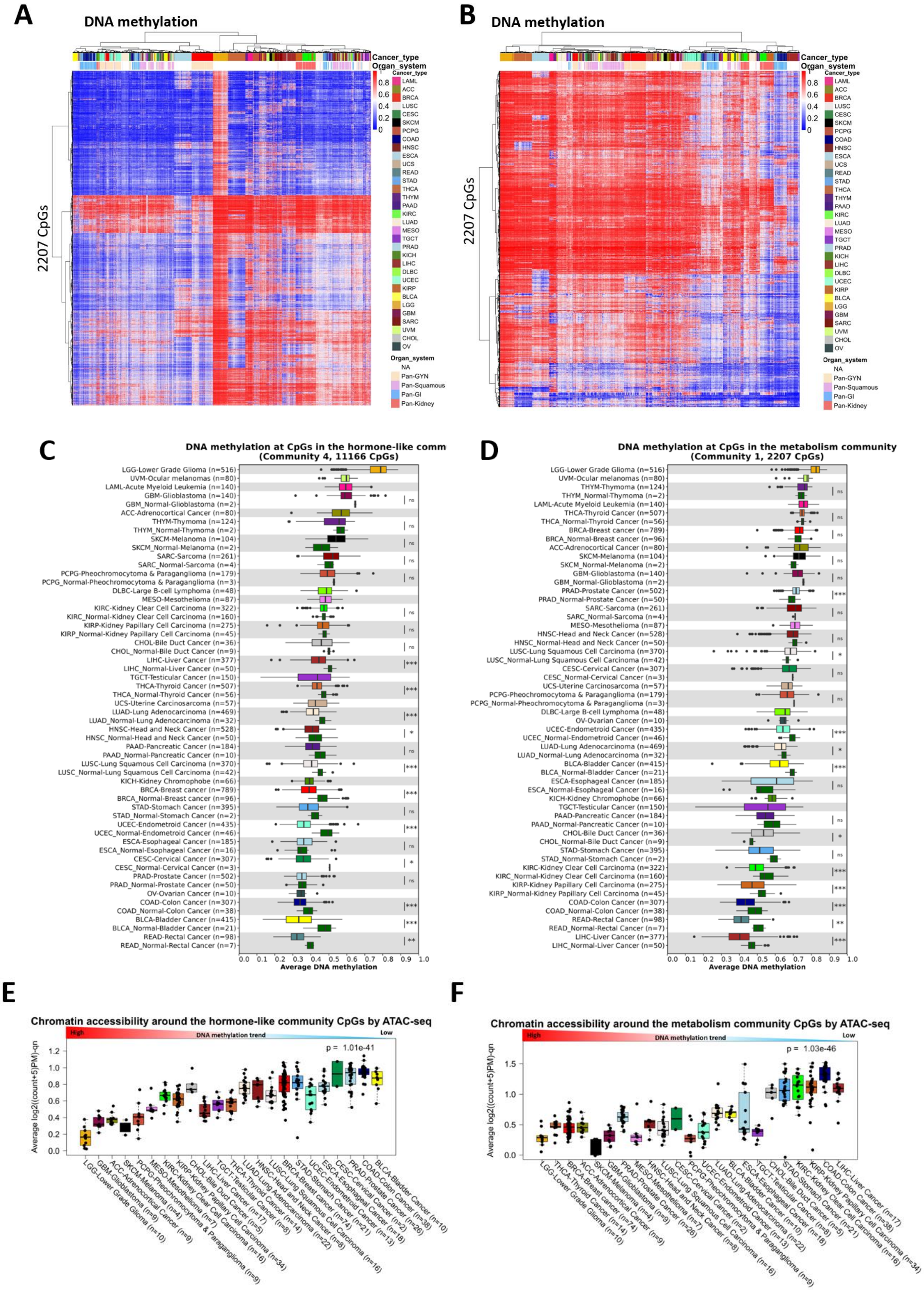
A
DNA methylation
2207 CpGs
Cancer_type
Organ_system
B
DNA methylation
2207 CpGs
Cancer_type
Organ_system
LAML
ACC
BRCA
LUSC
CESC
SKCM
PCPG
COAD
HNSC
ESCA
UCS
READ
STAD
THCA
THYM
PAAD
KIRC
LUAD
MESO
TGCT
PRAD
KICH
LIHC
DLBC
UCEC
KIRP
BLCA
LGG
GBM
SARC
UVM
CHOL
OV
NA
Pan-GYN
Pan-Squamous
Pan-GI
Pan-Kidney
C
DNA methylation at CpGs in the hormone-like comm
(Community 4, 11166 CpGs)
Average DNA methylation
D
DNA methylation at CpGs in the metabolism community
(Community 1, 2207 CpGs)
Average DNA methylation
E
Chromatin accessibility around the hormone-like community CpGs by ATAC-seq
High
DNA methylation trend
Low
p = 1.01e-41
Average log2((count+5)PM)-qn
F
Chromatin accessibility around the metabolism community CpGs by ATAC-seq
High
DNA methylation trend
Low
p = 1.03e-46
Average log2((count+5)PM)-qn

**Supplementary figure 6. DNA methylation at CpGs in the hormone-like and metabolism communities.** Heatmaps showing the DNA methylation levels of the hormone-like (A) and metabolism (B) communities CpGs in the TCGA-PANCAN dataset (n=8229) following unsupervised hierarchical clustering of DNA methylation levels. Rows represent CpGs and tumor samples represent the columns. Un-methylated and methylated CpGs are shown in blue and red points respectively. Histopathological features including cancer type and organ system are indicated in the columns. Box plots showing the average DNA methylation levels at the hormone-like (C) and metabolism (D) communities CpGs by TCGA cancer type. DNA methylation levels from available normal samples are included. BH-corrected Wilcoxon-test p-values are denoted. (C) Box plots showing the accessibility of the hormone-like (E) and metabolism (F) communities CpGs for different TCGA-PANCAN cancer types obtained by ATAC-seq. A higher value represents less compact chromatin and vice versa.

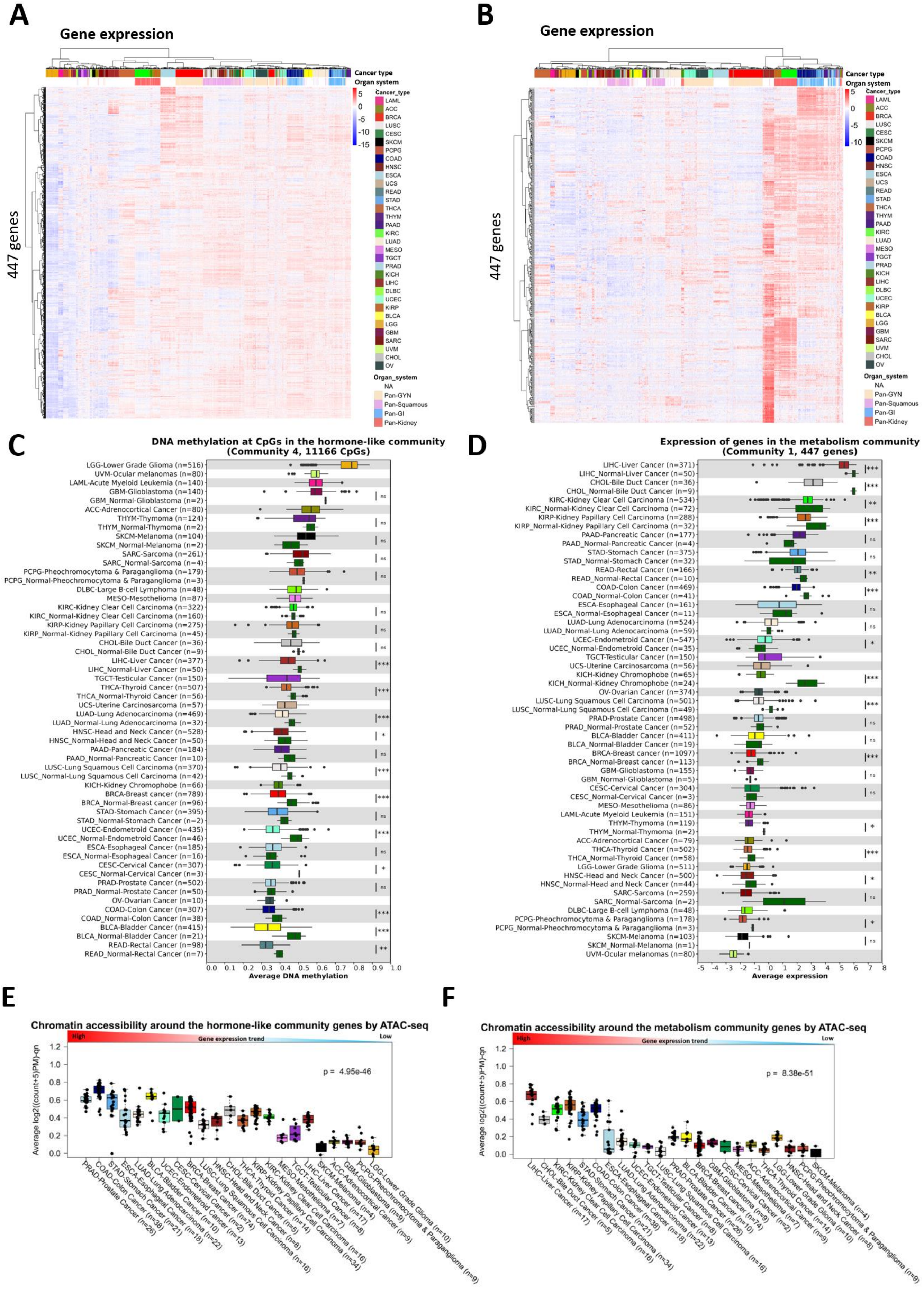

A
Gene expression
Cancer type
Organ system
447 genes
5
0
-5
-10
-15
Cancer_type
LAML
ACC
BRCA
LUSC
CESC
SKCM
PCPG
COAD
HNSC
ESCA
UCS
READ
STAD
THCA
THYM
PAAD
KIRC
LUAD
MESO
TGCT
PRAD
KICH
LIHC
DLBC
UCEC
KIRP
BLCA
LGG
GBM
SARC
UVM
CHOL
OV
Organ_system
NA
Pan-GYN
Pan-Squamous
Pan-GI
Pan-Kidney
B
Gene expression
Cancer type
Organ system
447 genes
5
0
-5
-10
C
DNA methylation at CpGs in the hormone-like community
(Community 4, 11166 CpGs)
LGG-Lower Grade Glioma (n=516)
UVM-Ocular melanomas (n=80)
LAML-Acute Myeloid Leukemia (n=140)
GBM-Glioblastoma (n=140)
GBM_Normal-Glioblastoma (n=2)
ACC-Adrenocortical Cancer (n=80)
THYM-Thymoma (n=124)
THYM_Normal-Thymoma (n=2)
SKCM-Melanoma (n=104)
SKCM_Normal-Melanoma (n=2)
SARC-Sarcoma (n=261)
SARC_Normal-Sarcoma (n=4)
PCPG-Pheochromocytoma & Paraganglioma (n=179)
PCPG_Normal-Pheochromocytoma & Paraganglioma (n=3)
DLBC-Large B-cell Lymphoma (n=48)
MESO-Mesothelioma (n=87)
KIRC-Kidney Clear Cell Carcinoma (n=322)
KIRC_Normal-Kidney Clear Cell Carcinoma (n=160)
KIRP-Kidney Papillary Cell Carcinoma (n=275)
KIRP_Normal-Kidney Papillary Cell Carcinoma (n=45)
CHOL-Bile Duct Cancer (n=36)
CHOL_Normal-Bile Duct Cancer (n=9)
LIHC-Liver Cancer (n=377)
LIHC_Normal-Liver Cancer (n=50)
TGCT-Testicular Cancer (n=150)
THCA-Thyroid Cancer (n=507)
THCA_Normal-Thyroid Cancer (n=56)
UCS-Uterine Carcinosarcoma (n=57)
LUAD-Lung Adenocarcinoma (n=469)
LUAD_Normal-Lung Adenocarcinoma (n=32)
HNSC-Head and Neck Cancer (n=528)
HNSC_Normal-Head and Neck Cancer (n=50)
PAAD-Pancreatic Cancer (n=184)
PAAD_Normal-Pancreatic Cancer (n=10)
LUSC-Lung Squamous Cell Carcinoma (n=370)
LUSC_Normal-Lung Squamous Cell Carcinoma (n=42)
KICH-Kidney Chromophobe (n=66)
BRCA-Breast cancer (n=789)
BRCA_Normal-Breast cancer (n=96)
STAD-Stomach Cancer (n=395)
STAD_Normal-Stomach Cancer (n=2)
UCEC-Endometroid Cancer (n=435)
UCEC_Normal-Endometroid Cancer (n=46)
ESCA-Esophageal Cancer (n=185)
ESCA_Normal-Esophageal Cancer (n=16)
CESC-Cervical Cancer (n=307)
CESC_Normal-Cervical Cancer (n=3)
PRAD-Prostate Cancer (n=502)
PRAD_Normal-Prostate Cancer (n=50)
OV-Ovarian Cancer (n=10)
COAD-Colon Cancer (n=307)
COAD_Normal-Colon Cancer (n=38)
BLCA-Bladder Cancer (n=415)
BLCA_Normal-Bladder Cancer (n=21)
READ-Rectal Cancer (n=98)
READ_Normal-Rectal Cancer (n=7)
ns
ns
ns
ns
ns
ns
ns
ns
***
***
***
*
ns
***
***
ns
***
*
ns
***
***
**
0.0 0.1 0.2 0.3 0.4 0.5 0.6 0.7 0.8 0.9 1.0
Average DNA methylation
D
Expression of genes in the metabolism community
(Community 1, 447 genes)
LIHC-Liver Cancer (n=371)
LIHC_Normal-Liver Cancer (n=50)
CHOL-Bile Duct Cancer (n=36)
CHOL_Normal-Bile Duct Cancer (n=9)
KIRC-Kidney Clear Cell Carcinoma (n=534)
KIRC_Normal-Kidney Clear Cell Carcinoma (n=72)
KIRP-Kidney Papillary Cell Carcinoma (n=288)
KIRP_Normal-Kidney Papillary Cell Carcinoma (n=32)
PAAD-Pancreatic Cancer (n=177)
PAAD_Normal-Pancreatic Cancer (n=4)
STAD-Stomach Cancer (n=375)
STAD_Normal-Stomach Cancer (n=32)
READ-Rectal Cancer (n=166)
READ_Normal-Rectal Cancer (n=10)
COAD-Colon Cancer (n=469)
COAD_Normal-Colon Cancer (n=41)
ESCA-Esophageal Cancer (n=161)
ESCA_Normal-Esophageal Cancer (n=11)
LUAD-Lung Adenocarcinoma (n=524)
LUAD_Normal-Lung Adenocarcinoma (n=59)
UCEC-Endometroid Cancer (n=547)
UCEC_Normal-Endometroid Cancer (n=35)
TGCT-Testicular Cancer (n=150)
UCS-Uterine Carcinosarcoma (n=56)
KICH-Kidney Chromophobe (n=65)
KICH_Normal-Kidney Chromophobe (n=24)
OV-Ovarian Cancer (n=374)
LUSC-Lung Squamous Cell Carcinoma (n=501)
LUSC_Normal-Lung Squamous Cell Carcinoma (n=49)
PRAD-Prostate Cancer (n=498)
PRAD_Normal-Prostate Cancer (n=52)
BLCA-Bladder Cancer (n=411)
BLCA_Normal-Bladder Cancer (n=19)
BRCA-Breast cancer (n=1097)
BRCA_Normal-Breast cancer (n=113)
GBM-Glioblastoma (n=155)
GBM_Normal-Glioblastoma (n=5)
CESC-Cervical Cancer (n=304)
CESC_Normal-Cervical Cancer (n=3)
MESO-Mesothelioma (n=86)
LAML-Acute Myeloid Leukemia (n=151)
THYM-Thymoma (n=119)
THYM_Normal-Thymoma (n=2)
ACC-Adrenocortical Cancer (n=79)
THCA-Thyroid Cancer (n=502)
THCA_Normal-Thyroid Cancer (n=58)
LGG-Lower Grade Glioma (n=511)
HNSC-Head and Neck Cancer (n=500)
HNSC_Normal-Head and Neck Cancer (n=44)
SARC-Sarcoma (n=259)
SARC_Normal-Sarcoma (n=2)
DLBC-Large B-cell Lymphoma (n=48)
PCPG-Pheochromocytoma & Paraganglioma (n=178)
PCPG_Normal-Pheochromocytoma & Paraganglioma (n=3)
SKCM-Melanoma (n=103)
SKCM_Normal-Melanoma (n=1)
UVM-Ocular melanomas (n=80)
***
***
**
***
ns
ns
**
***
ns
ns
*
***
***
ns
ns
***
ns
ns
*
***
*
ns
*
ns
-5 -4 -3 -2 -1 0 1 2 3 4 5 6 7 8
Average expression
E
Chromatin accessibility around the hormone-like community genes by ATAC-seq
High
Gene expression trend
Low
Average log2((count+5)PM)-qn
1.2
1.0
0.8
0.6
0.4
0.2
0.0
p = 4.95e-46
PRAD-Prostate Cancer (n=26)
COAD-Colon Cancer (n=38)
STAD-Stomach Cancer (n=21)
ESCA-Esophageal Cancer (n=18)
LUAD-Lung Adenocarcinoma (n=22)
BLCA-Bladder Cancer (n=10)
UCEC-Endometroid Cancer (n=13)
CESC-Cervical Cancer (n=2)
BRCA-Breast cancer (n=74)
HNSC-Head and Neck Cancer (n=9)
LUSC-Lung Squamous Cell Carcinoma (n=16)
CHOL-Bile Duct Cancer (n=5)
THCA-Thyroid Cancer (n=14)
KIRP-Kidney Papillary Cell Carcinoma (n=34)
KIRC-Kidney Clear Cell Carcinoma (n=16)
MESO-Mesothelioma (n=7)
TGCT-Testicular Cancer (n=8)
LIHC-Liver Cancer (n=17)
SKCM-Melanoma (n=4)
ACC-Adrenocortical Cancer (n=9)
GBM-Glioblastoma (n=9)
PCPG-Pheochromocytoma & Paraganglioma (n=9)
LGG-Lower Grade Glioma (n=10)
F
Chromatin accessibility around the metabolism community genes by ATAC-seq
High
Gene expression trend
Low
Average log2((count+5)PM)-qn
1.2
1.0
0.8
0.6
0.4
0.2
0.0
p = 8.38e-51
LIHC-Liver Cancer (n=17)
CHOL-Bile Duct Cancer (n=5)
KIRC-Kidney Clear Cell Carcinoma (n=16)
KIRP-Kidney Papillary Cell Carcinoma (n=34)
STAD-Stomach Cancer (n=21)
COAD-Colon Cancer (n=38)
ESCA-Esophageal Cancer (n=18)
LUAD-Lung Adenocarcinoma (n=22)
UCEC-Endometroid Cancer (n=13)
TGCT-Testicular Cancer (n=8)
LUSC-Lung Squamous Cell Carcinoma (n=16)
PRAD-Prostate Cancer (n=26)
BLCA-Bladder Cancer (n=10)
BRCA-Breast cancer (n=74)
GBM-Glioblastoma (n=9)
CESC-Cervical Cancer (n=2)
MESO-Mesothelioma (n=7)
ACC-Adrenocortical Cancer (n=9)
THCA-Thyroid Cancer (n=14)
LGG-Lower Grade Glioma (n=10)
HNSC-Head and Neck Cancer (n=9)
PCPG-Pheochromocytoma & Paraganglioma (n=9)
SKCM-Melanoma (n=4)

**Supplementary figure 7. Expression of genes in the hormone-like and metabolism communities.** Hierarchical clustering of the expression levels of the genes in the hormone-like (A) and metabolism (B) communities in the TCGA-PANCAN dataset (n=9875). Rows represent genes and column tumor samples. Red points indicate high expression and blue points indicate low expression. Histopathological features including organ system and cancer type are included. Box plots showing the average expression levels of the hormone-like (C) and metabolism (D) community genes by cancer type. Tumor samples with normal tissue are included. BH-corrected Wilcoxon-test p-values are denoted. Chromatin accessibility around the hormone-like (E) and metabolism (F) community genes in the TCGA-PANCAN tumor samples.

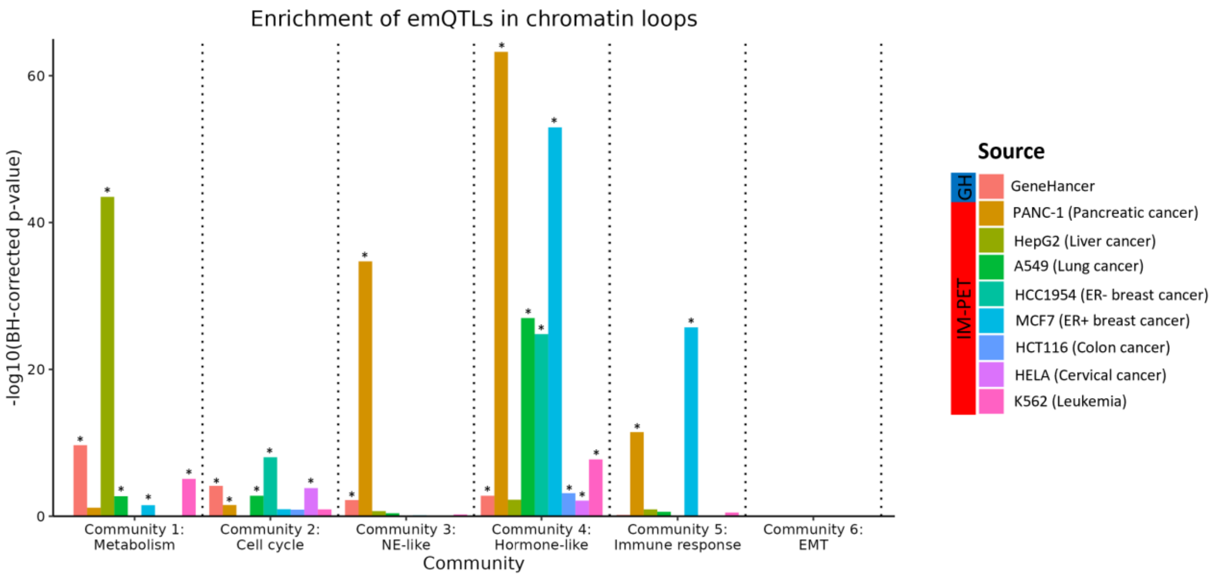


**Supplementary figure 8. Enrichment of emQTL in chromatin loops.** Bar plot showing the enrichment of pan-cancer emQTL from each emQTL community in GeneHancer and IM-PET loops (PANC-1, HepG2, A549, HCC1954, MCF7, HCT116, HELA, and K562). The height of the bars represents the -log10(BH-corrected p-values) obtained by hypergeometric testing. Statistically significant enrichments are marked with an asterisk (BH-corrected $p\text{-value}<0.05$).

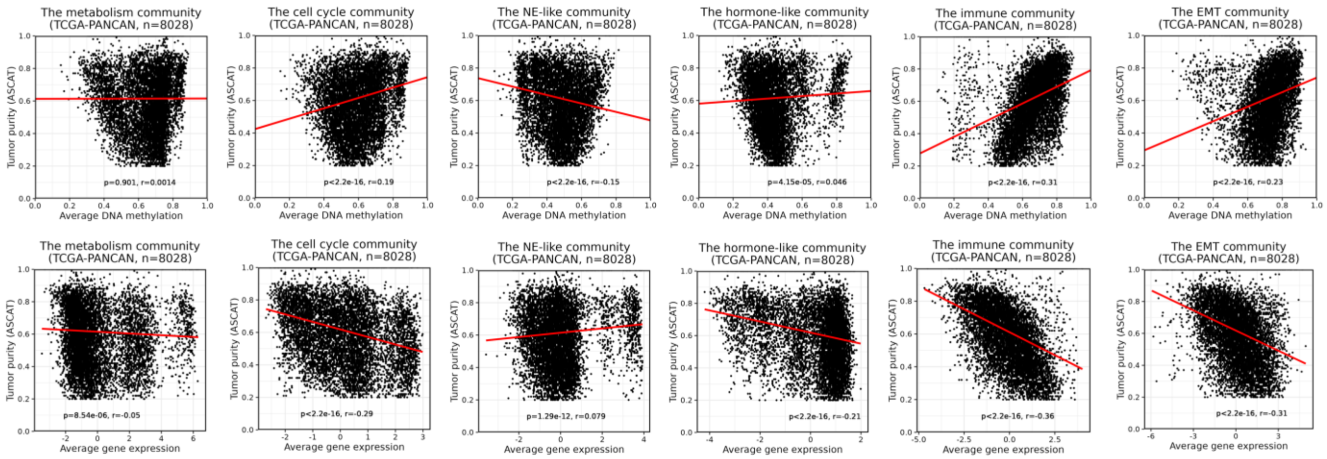


**Supplementary figure 9. Association between DNA methylation and expression versus tumor purity.** Scatter plots showing the correlation between DNA methylation at the emQTL-CpGs and expression of emQTL genes versus the ASCAT tumor purity estimates in the TCGA-PANCAN dataset. P-values and Pearson correlation coefficients are denoted.

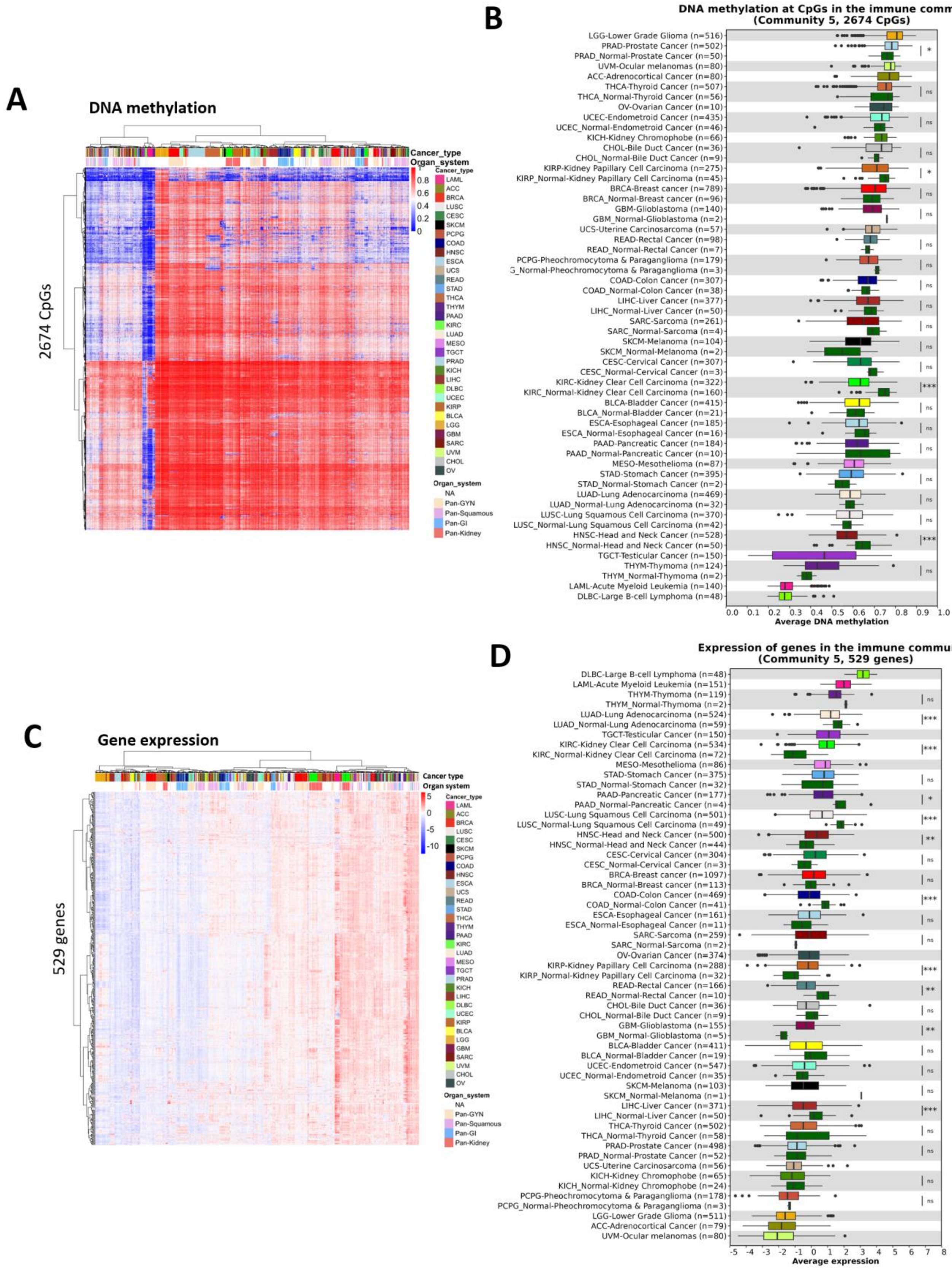


**Supplementary figure 10. DNA methylation at CpGs and expression of genes in the immune community.** (A) Unsupervised hierarchical clustering of DNA methylation levels at the CpGs in the immune community in TCGA (n=8229). The red points indicate methylated CpGs and the blue points indicate unmethylated. Rows represent CpGs and columns represent tumor samples. (B) Box plot showing the average DNA methylation levels at the immune community CpGs by TCGA cancer type. DNA methylation levels from available normal samples are included. BH-corrected Wilcoxon-test p-values are denoted. (C) Unsupervised hierarchical clustering of gene expression levels of the immune community genes in each TCGA-PANCAN dataset (n=9875). Rows represent genes and columns represent tumor samples.

Red points indicate high expression and blue points low expression. Histopathological features including organ system and cancer type are included. (D) Box plot showing the average expression levels of the immune community genes by cancer types. Tumor samples with normal tissue are included. BH-corrected Wilcoxon-test p-values are denoted.

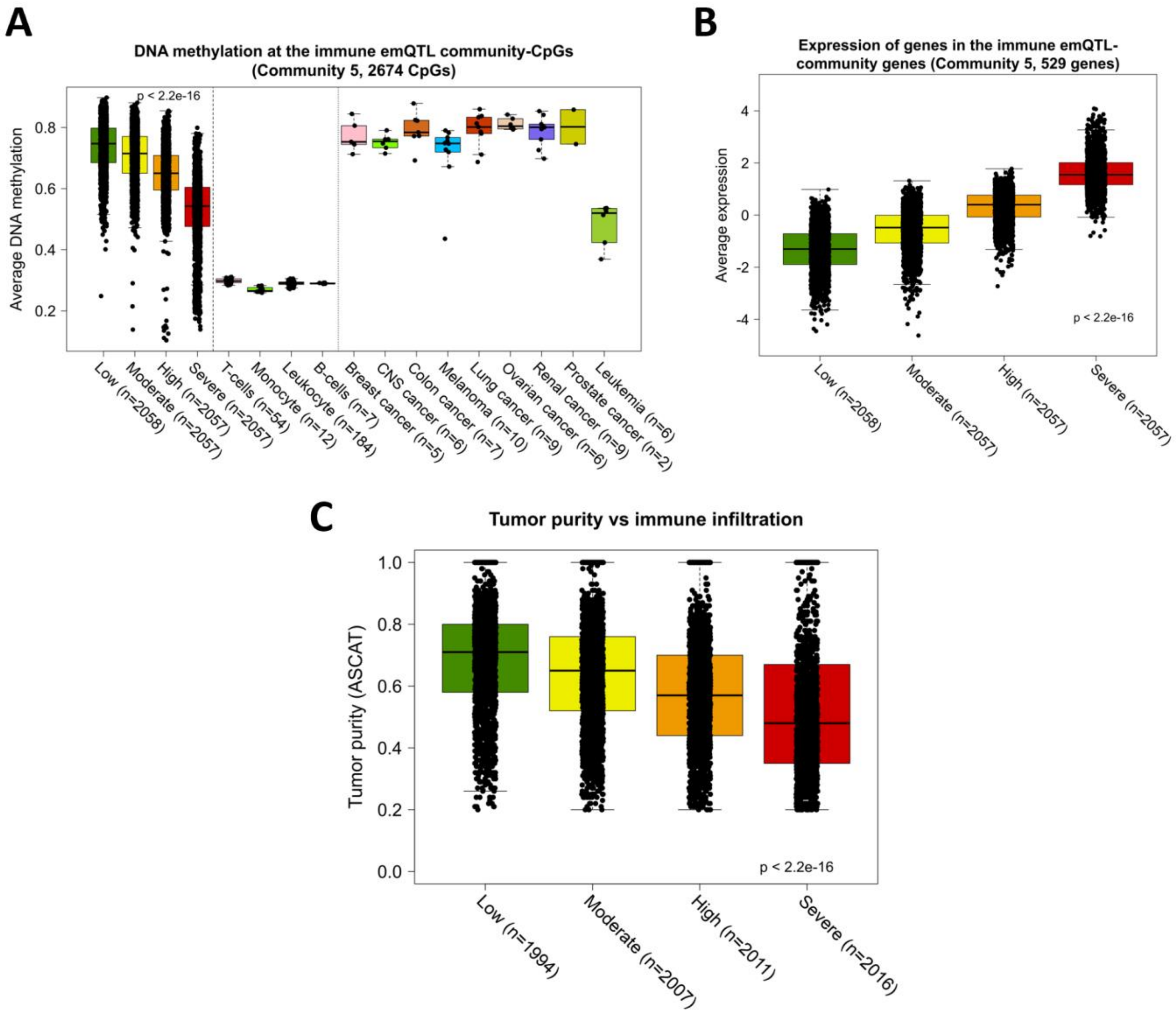


**Supplementary figure 11. emQTL community 5 is linked to immune infiltration.** (A) Box plot showing the link between DNA methylation at the immune community CpGs and immune cell infiltration. The level of immune infiltration in tumor samples from TCGA was determined using the xCell deconvolution tool [49]. Tumor samples were divided into quartile groups based on the severity of immune infiltration; low, moderate, high, and severe. DNA methylation levels at the immune community CpGs in isolated immune cells (T-cells, monocytes, B-cells, and Leukocytes) and cancer cell lines from the NCI-60 cancer cell lines are included. (B) Expression of the immune community genes in relation to the xCell-derived immune score. (C) Box plot showing the association between the xCell-derived immune score and tumor purity estimates obtained by ASCAT. Kruskal-Wallis test p-values determined by comparing the quartile groups are denoted in each box plot (A-C).

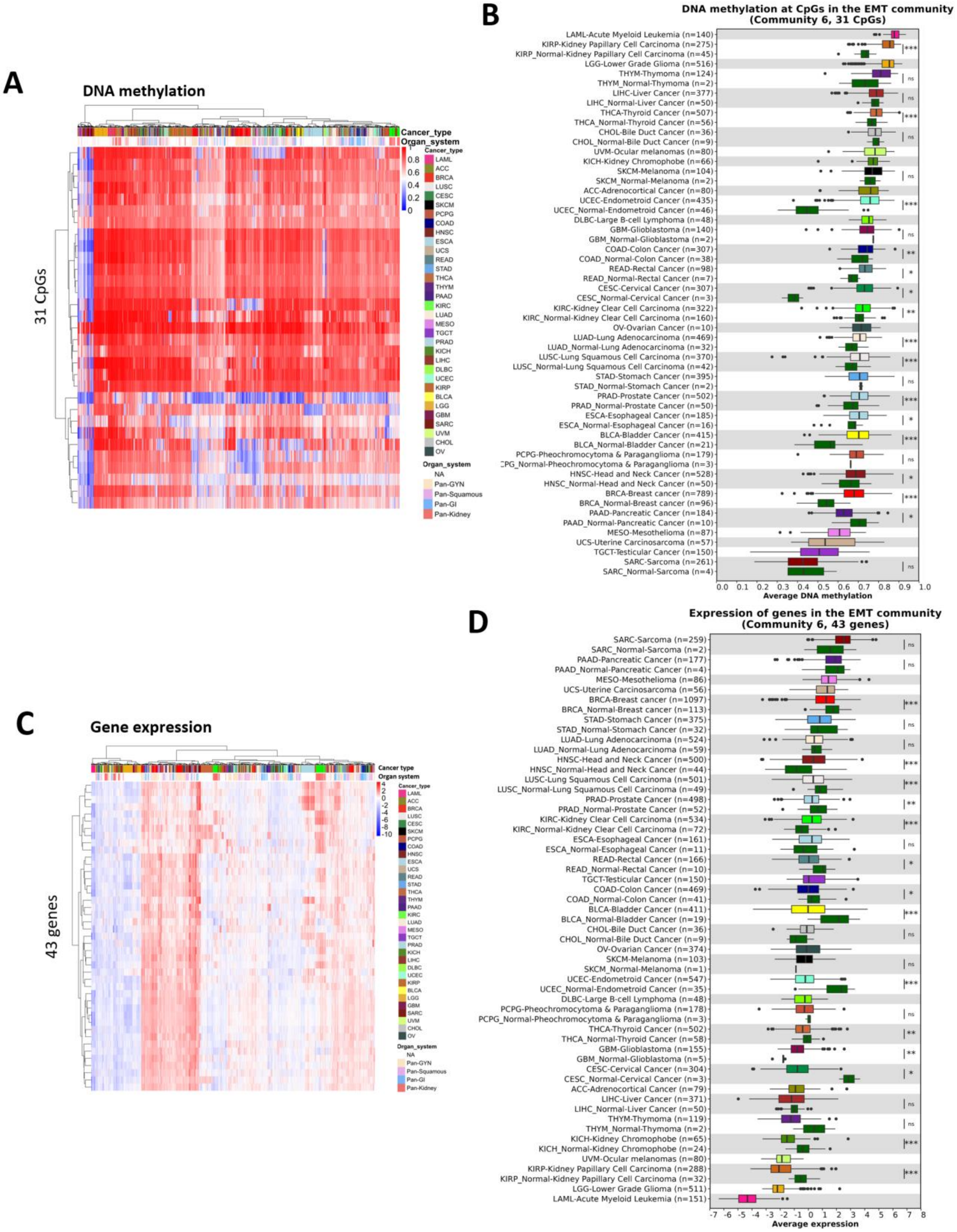


**Supplementary figure 12. DNA methylation at CpGs and expression of genes in the EMT community.** (A) Unsupervised hierarchical clustering of DNA methylation levels at the EMT community CpGs in TCGA-PANCAN (n=8229). Red points indicate methylated CpGs and blue points indicate unmethylated. Rows represent CpGs and columns represent tumor samples. (B) Box plot showing the average DNA methylation levels at the EMT community CpGs by cancer type. DNA methylation levels from available normal samples are included. BH-corrected Wilcoxon-test p-values are denoted. (C) Unsupervised hierarchical clustering of gene expression levels of the EMT community genes in each TCGA-PANCAN dataset (n=9875). Rows represent genes and column tumor samples. Red points indicate high expression and blue points indicate low expression. Histopathological features including organ system and cancer type are included. (D) Box plot showing the average expression levels of the EMT community genes by cancer types. Tumor samples with normal tissue are included. BH-corrected Wilcoxon-test p-values are denoted.

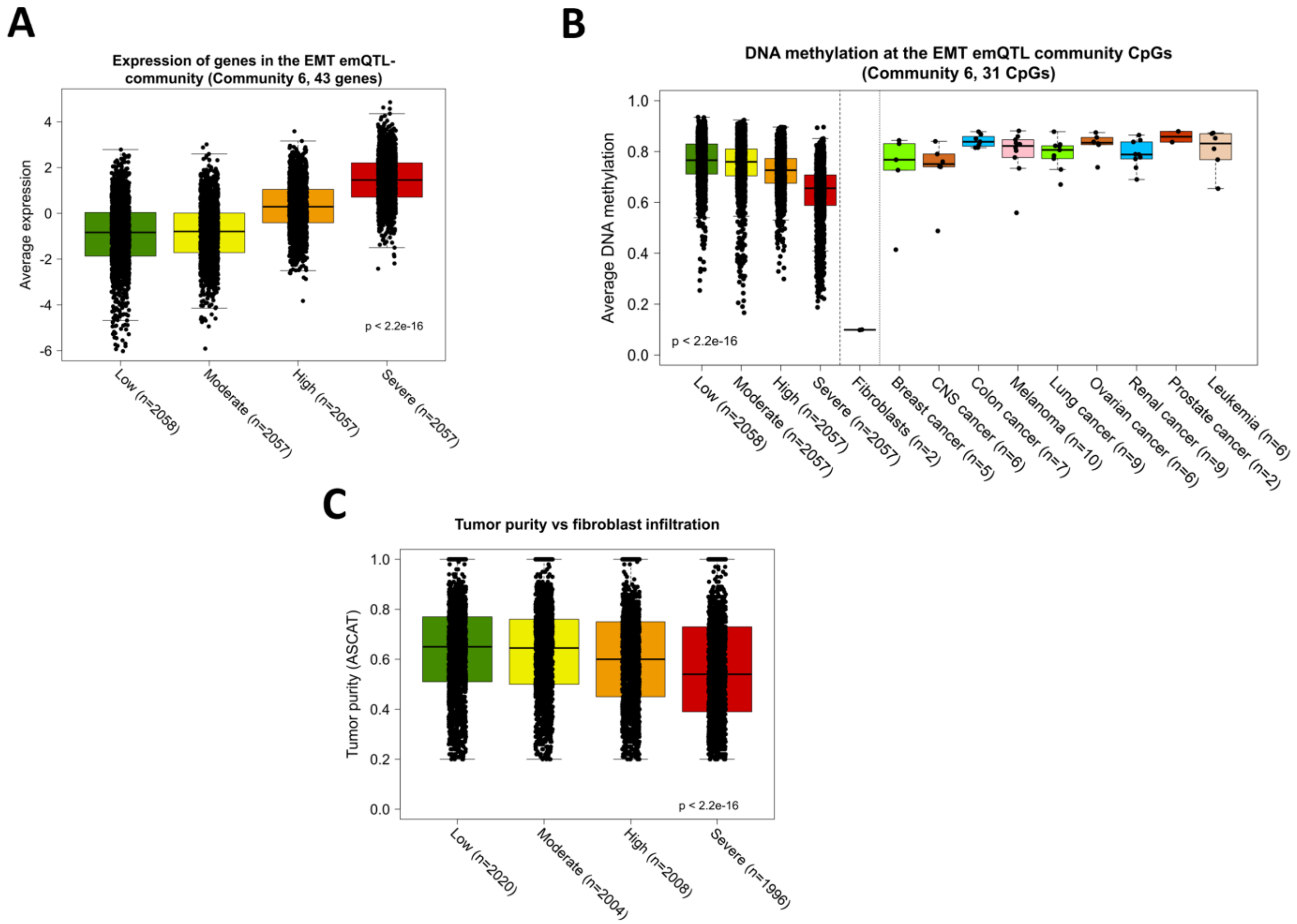


**Supplementary figure 13. emQTL community 6 is associated with fibroblast infiltration.** (A) Box plot showing the association between expression of the EMT community genes in relation to fibroblast infiltration. Fibroblast infiltration level in the tumor samples in TCGA was determined by the xCell deconvolution tool. Tumor samples were divided into quartile groups based on the severity of fibroblast infiltration; low, moderate, high, and severe. (B) The association between fibroblast infiltration and DNA methylation at the EMT community CpGs. DNA methylation levels obtained from human mammary fibroblasts and the NCI-60 cancer cell lines are also included. (C) shows the association between fibroblast infiltration and tumor purity estimates from ASCAT. Kruskal-Wallis test p-values comparing the quartile groups are denoted in each plot.

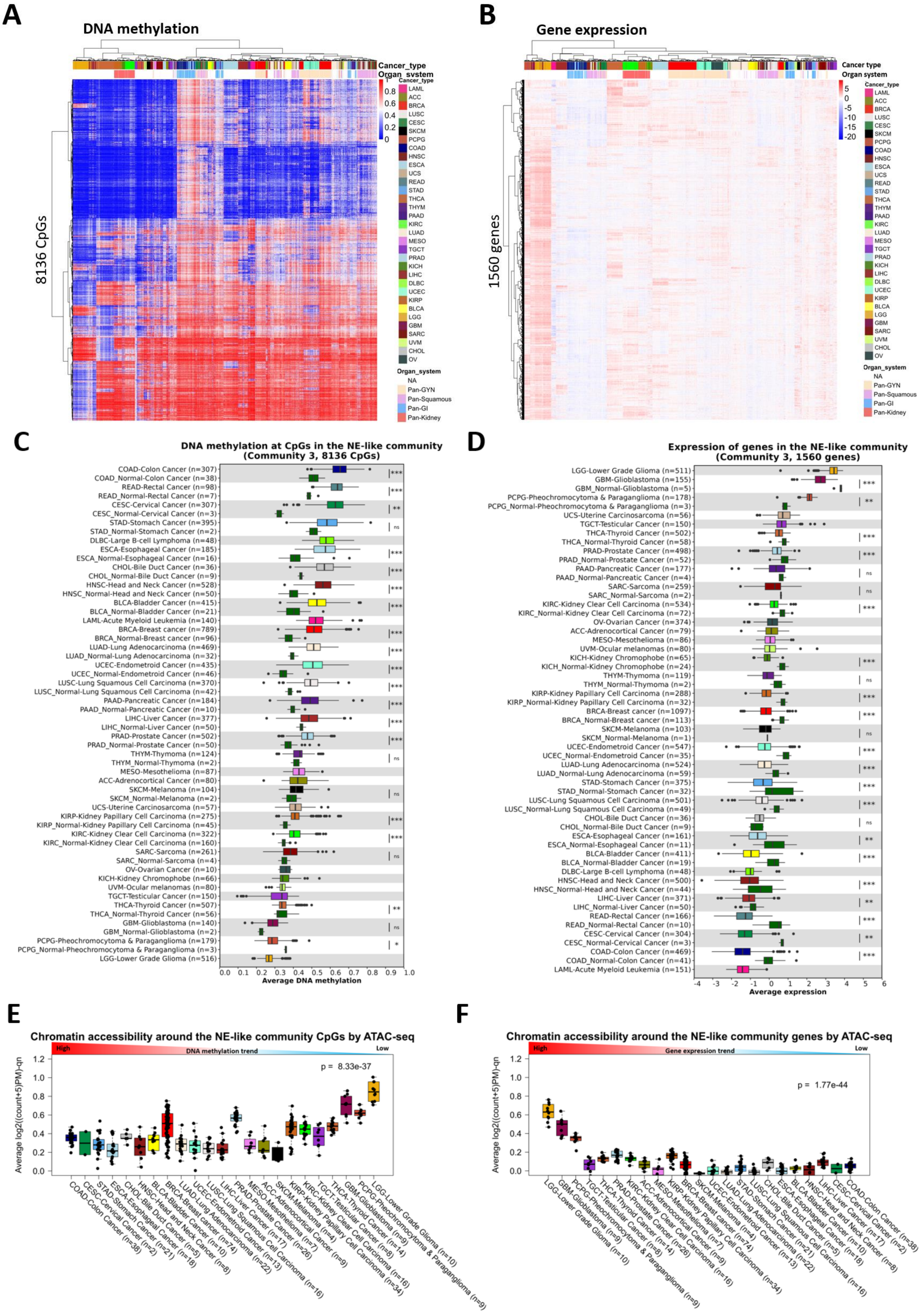
A
DNA methylation
Cancer_type
Organ_system
8136 CpGs
1
0.8
0.6
0.4
0.2
0
Cancer_type
LAML
ACC
BRCA
LUSC
CESC
SKCM
PCPG
COAD
HNSC
ESCA
UCS
READ
STAD
THCA
THYM
PAAD
KIRC
LUAD
MESO
TGCT
PRAD
KICH
LIHC
DLBC
UCEC
KIRP
BLCA
LGG
GBM
SARC
UVM
CHOL
OV
Organ_system
NA
Pan-GYN
Pan-Squamous
Pan-GI
Pan-Kidney
B
Gene expression
Cancer type
Organ system
1560 genes
5
0
-5
-10
-15
-20
Cancer_type
LAML
ACC
BRCA
LUSC
CESC
SKCM
PCPG
COAD
HNSC
ESCA
UCS
READ
STAD
THCA
THYM
PAAD
KIRC
LUAD
MESO
TGCT
PRAD
KICH
LIHC
DLBC
UCEC
KIRP
BLCA
LGG
GBM
SARC
UVM
CHOL
OV
Organ_system
NA
Pan-GYN
Pan-Squamous
Pan-GI
Pan-Kidney
C
DNA methylation at CpGs in the NE-like community
(Community 3, 8136 CpGs)
COAD-Colon Cancer (n=307)
COAD_Normal-Colon Cancer (n=38)
READ-Rectal Cancer (n=98)
READ_Normal-Rectal Cancer (n=7)
CESC-Cervical Cancer (n=307)
CESC_Normal-Cervical Cancer (n=3)
STAD-Stomach Cancer (n=395)
STAD_Normal-Stomach Cancer (n=2)
DLBC-Large B-cell Lymphoma (n=48)
ESCA-Esophageal Cancer (n=185)
ESCA_Normal-Esophageal Cancer (n=16)
CHOL-Bile Duct Cancer (n=36)
CHOL_Normal-Bile Duct Cancer (n=9)
HNSC-Head and Neck Cancer (n=528)
HNSC_Normal-Head and Neck Cancer (n=50)
BLCA-Bladder Cancer (n=415)
BLCA_Normal-Bladder Cancer (n=21)
LAML-Acute Myeloid Leukemia (n=140)
BRCA-Breast cancer (n=789)
BRCA_Normal-Breast cancer (n=96)
LUAD-Lung Adenocarcinoma (n=469)
LUAD_Normal-Lung Adenocarcinoma (n=32)
UCEC-Endometroid Cancer (n=435)
UCEC_Normal-Endometroid Cancer (n=46)
LUSC-Lung Squamous Cell Carcinoma (n=370)
LUSC_Normal-Lung Squamous Cell Carcinoma (n=42)
PAAD-Pancreatic Cancer (n=184)
PAAD_Normal-Pancreatic Cancer (n=10)
LIHC-Liver Cancer (n=377)
LIHC_Normal-Liver Cancer (n=50)
PRAD-Prostate Cancer (n=502)
PRAD_Normal-Prostate Cancer (n=50)
THYM-Thymoma (n=124)
THYM_Normal-Thymoma (n=2)
MESO-Mesothelioma (n=87)
ACC-Adrenocortical Cancer (n=80)
SKCM-Melanoma (n=104)
SKCM_Normal-Melanoma (n=2)
UCS-Uterine Carcinosarcoma (n=57)
KIRP-Kidney Papillary Cell Carcinoma (n=275)
KIRP_Normal-Kidney Papillary Cell Carcinoma (n=45)
KIRC-Kidney Clear Cell Carcinoma (n=322)
KIRC_Normal-Kidney Clear Cell Carcinoma (n=160)
SARC-Sarcoma (n=261)
SARC_Normal-Sarcoma (n=4)
OV-Ovarian Cancer (n=10)
KICH-Kidney Chromophobe (n=66)
UVM-Ocular melanomas (n=80)
TGCT-Testicular Cancer (n=150)
THCA-Thyroid Cancer (n=507)
THCA_Normal-Thyroid Cancer (n=56)
GBM-Glioblastoma (n=140)
GBM_Normal-Glioblastoma (n=2)
PCPG-Pheochromocytoma & Paraganglioma (n=179)
PCPG_Normal-Pheochromocytoma & Paraganglioma (n=3)
LGG-Lower Grade Glioma (n=516)
***
***
**
ns
***
***
***
***
***
***
***
***
***
***
***
ns
ns
***
***
ns
**
ns
*
0.0 0.1 0.2 0.3 0.4 0.5 0.6 0.7 0.8 0.9 1.0
Average DNA methylation
D
Expression of genes in the NE-like community
(Community 3, 1560 genes)
LGG-Lower Grade Glioma (n=511)
GBM-Glioblastoma (n=155)
GBM_Normal-Glioblastoma (n=5)
PCPG-Pheochromocytoma & Paraganglioma (n=178)
PCPG_Normal-Pheochromocytoma & Paraganglioma (n=3)
UCS-Uterine Carcinosarcoma (n=56)
TGCT-Testicular Cancer (n=150)
THCA-Thyroid Cancer (n=502)
THCA_Normal-Thyroid Cancer (n=58)
PRAD-Prostate Cancer (n=498)
PRAD_Normal-Prostate Cancer (n=52)
PAAD-Pancreatic Cancer (n=177)
PAAD_Normal-Pancreatic Cancer (n=4)
SARC-Sarcoma (n=259)
SARC_Normal-Sarcoma (n=2)
KIRC-Kidney Clear Cell Carcinoma (n=534)
KIRC_Normal-Kidney Clear Cell Carcinoma (n=72)
OV-Ovarian Cancer (n=374)
ACC-Adrenocortical Cancer (n=79)
MESO-Mesothelioma (n=86)
UVM-Ocular melanomas (n=80)
KICH-Kidney Chromophobe (n=65)
KICH_Normal-Kidney Chromophobe (n=24)
THYM-Thymoma (n=119)
THYM_Normal-Thymoma (n=2)
KIRP-Kidney Papillary Cell Carcinoma (n=288)
KIRP_Normal-Kidney Papillary Cell Carcinoma (n=32)
BRCA-Breast cancer (n=1097)
BRCA_Normal-Breast cancer (n=113)
SKCM-Melanoma (n=103)
SKCM_Normal-Melanoma (n=1)
UCEC-Endometroid Cancer (n=547)
UCEC_Normal-Endometroid Cancer (n=35)
LUAD-Lung Adenocarcinoma (n=524)
LUAD_Normal-Lung Adenocarcinoma (n=59)
STAD-Stomach Cancer (n=375)
STAD_Normal-Stomach Cancer (n=32)
LUSC-Lung Squamous Cell Carcinoma (n=501)
LUSC_Normal-Lung Squamous Cell Carcinoma (n=49)
CHOL-Bile Duct Cancer (n=36)
CHOL_Normal-Bile Duct Cancer (n=9)
ESCA-Esophageal Cancer (n=161)
ESCA_Normal-Esophageal Cancer (n=11)
BLCA-Bladder Cancer (n=411)
BLCA_Normal-Bladder Cancer (n=19)
DLBC-Large B-cell Lymphoma (n=48)
HNSC-Head and Neck Cancer (n=500)
HNSC_Normal-Head and Neck Cancer (n=44)
LIHC-Liver Cancer (n=371)
LIHC_Normal-Liver Cancer (n=50)
READ-Rectal Cancer (n=166)
READ_Normal-Rectal Cancer (n=10)
CESC-Cervical Cancer (n=304)
CESC_Normal-Cervical Cancer (n=3)
COAD-Colon Cancer (n=469)
COAD_Normal-Colon Cancer (n=41)
LAML-Acute Myeloid Leukemia (n=151)
***
**
***
***
ns
ns
***
***
ns
***
***
ns
***
***
***
***
ns
**
***
***
**
***
**
***
-4 -3 -2 -1 0 1 2 3 4 5 6
Average expression
E
Chromatin accessibility around the NE-like community CpGs by ATAC-seq
High
DNA methylation trend
Low
p = 8.33e-37
Average log2((count+5)PM)-qn
1.2
1.0
0.8
0.6
0.4
0.2
0.0
COAD-Colon Cancer (n=38)
CESC-Cervical Cancer (n=2)
STAD-Stomach Cancer (n=21)
ESCA-Esophageal Cancer (n=18)
CHOL-Bile Duct Cancer (n=5)
HNSC-Head and Neck Cancer (n=8)
BLCA-Bladder Cancer (n=10)
BRCA-Breast cancer (n=74)
LUAD-Lung Adenocarcinoma (n=22)
UCEC-Endometroid Cancer (n=13)
LUSC-Lung Squamous Cell Carcinoma (n=16)
LIHC-Liver Cancer (n=17)
PRAD-Prostate Cancer (n=26)
MESO-Mesothelioma (n=7)
ACC-Adrenocortical Cancer (n=9)
SKCM-Melanoma (n=4)
KIRP-Kidney Papillary Cell Carcinoma (n=34)
KIRC-Kidney Clear Cell Carcinoma (n=16)
TGCT-Testicular Cancer (n=8)
THCA-Thyroid Cancer (n=14)
GBM-Glioblastoma (n=9)
PCPG-Pheochromocytoma & Paraganglioma (n=9)
LGG-Lower Grade Glioma (n=10)
F
Chromatin accessibility around the NE-like community genes by ATAC-seq
High
Gene expression trend
Low
p = 1.77e-44
Average log2((count+5)PM)-qn
1.2
1.0
0.8
0.6
0.4
0.2
0.0
LGG-Lower Grade Glioma (n=10)
GBM-Glioblastoma (n=9)
PCPG-Pheochromocytoma & Paraganglioma (n=9)
TGCT-Testicular Cancer (n=8)
THCA-Thyroid Cancer (n=14)
PRAD-Prostate Cancer (n=26)
KIRC-Kidney Clear Cell Carcinoma (n=16)
ACC-Adrenocortical Cancer (n=9)
MESO-Mesothelioma (n=7)
KIRP-Kidney Papillary Cell Carcinoma (n=34)
BRCA-Breast cancer (n=74)
SKCM-Melanoma (n=4)
UCEC-Endometroid Cancer (n=13)
LUAD-Lung Adenocarcinoma (n=22)
STAD-Stomach Cancer (n=21)
LUSC-Lung Squamous Cell Carcinoma (n=16)
CHOL-Bile Duct Cancer (n=5)
ESCA-Esophageal Cancer (n=18)
BLCA-Bladder Cancer (n=10)
HNSC-Head and Neck Cancer (n=8)
LIHC-Liver Cancer (n=17)
CESC-Cervical Cancer (n=2)
COAD-Colon Cancer (n=38)

**Supplementary figure 14. DNA methylation at CpGs and expression of genes in the NE-like community.** (A) Hierarchical clustering of DNA methylation levels at the NE-like community CpGs in the TCGA-PANCAN dataset (n=8229). Rows represent CpGs and columns represent tumor samples. Methylated and unmethylated CpGs are shown as red and blue dots respectively. Cancer type and organ system of the cancer type are indicated. (B) Heatmap showing the expression levels of the NE-community genes in the TCGA-PANCAN dataset (n=9875). Rows and columns represent genes and samples respectively and were ordered by unsupervised hierarchical clustering. Blue points indicate low expression while red points indicate high. The samples are annotated by cancer type and organ system. (C) Box plot showing the average DNA methylation levels at the NE-like community CpGs by cancer type. DNA methylation levels from normal samples are included for those cancer types with data available. BH-corrected Wilcoxon-test p-values are denoted. (D) Box plot showing the average expression of genes in the NE-like community by cancer type. BH-corrected Wilcoxon test p-values are denoted by comparing expression levels of genes between tumor and normal samples. (E) Boxplot showing the chromatin accessibility around the CpGs in the NE-like community as determined by ATAC-seq of the tumor samples from TCGA-PANCAN. A high value represents more open chromatin while a low value represents more compact chromatin. The chromatin accessibility around the NE-like community genes is visualized in (F) for the TCGA-PANCAN dataset.

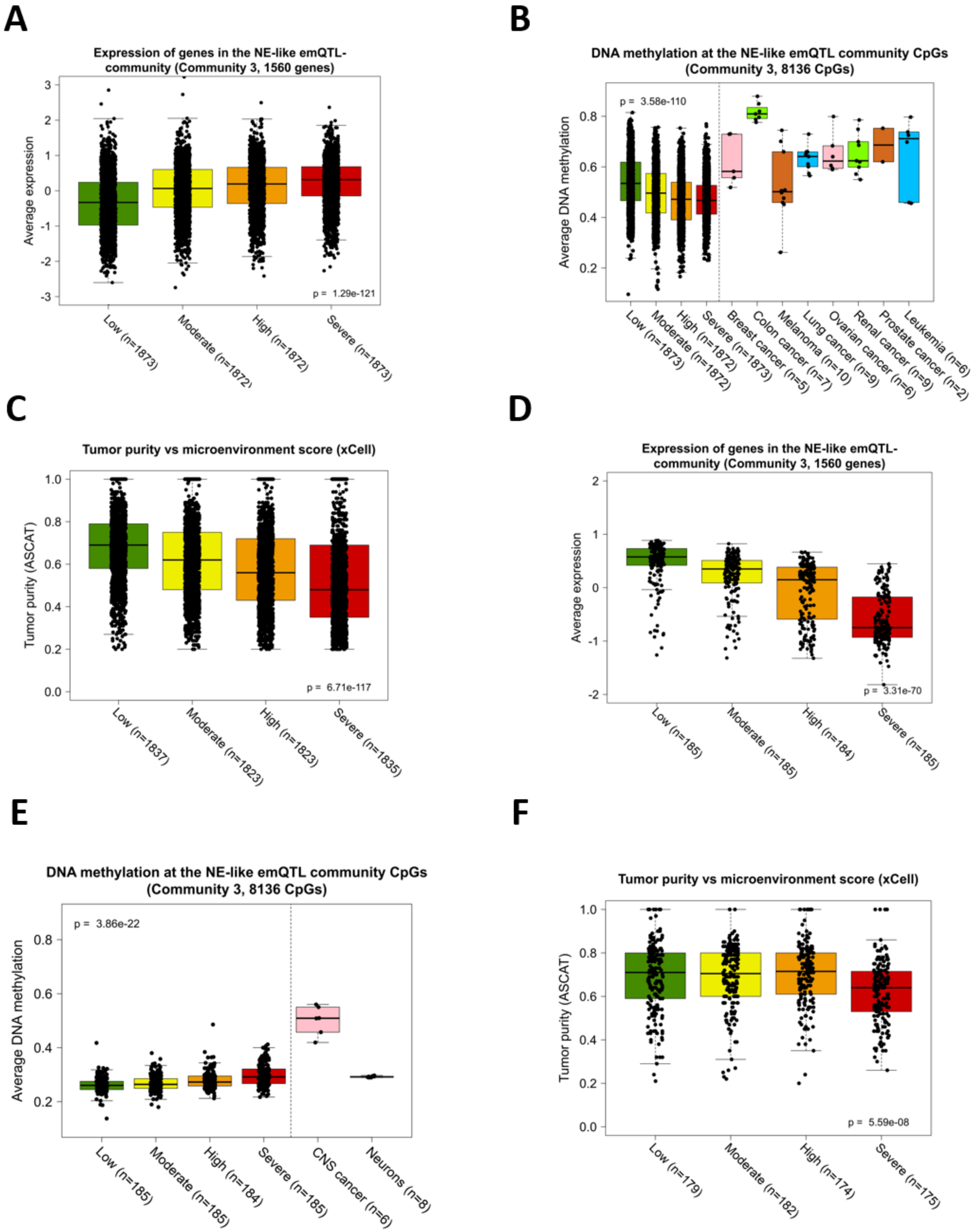


**Supplementary figure 15. Community 3 is linked to the infiltration of non-tumor cells.** Boxplots showing the expression of the NE-like community genes in the non-neuron (NN) related cancer types (A) and the neuron-related cancer types (LGG, PCPG, GBM; D) in TCGA. The tumors were divided into four quartile groups (Low, moderate, high, and severe) based on the severity of infiltration using the microenvironment score obtained from xCell. The DNA methylation levels at the NE-like community CpGs in NN and non-NN tumors are shown in (B) and (E) respectively. (C) and (F) shows the associations between the microenvironment score and the ASCAT tumor purity score in the non-NN and NN tumors respectively. Kruskal-Wallis test p-values are denoted in each box plot.

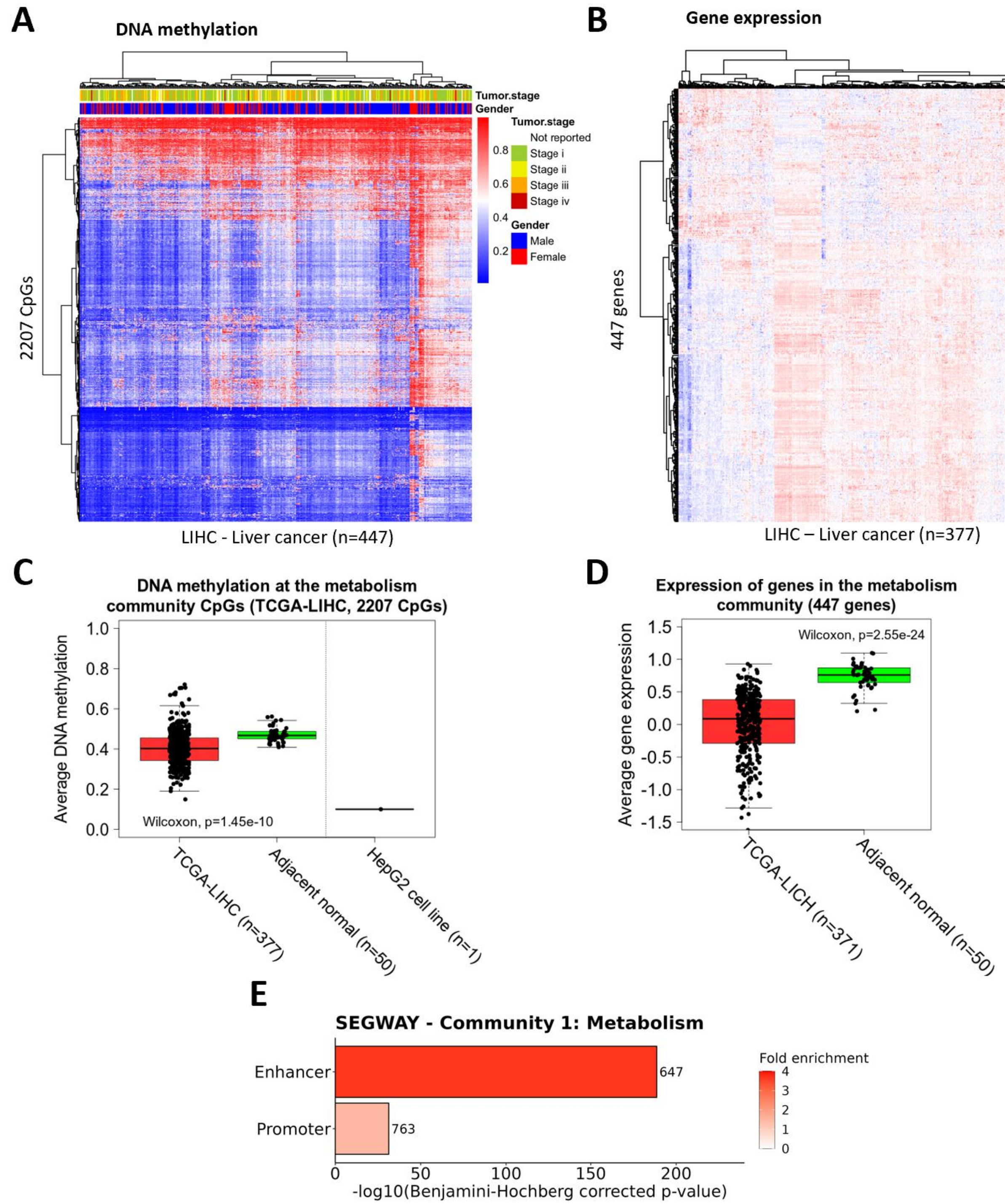


**Supplementary figure 16. DNA methylation at CpGs and expression of genes in the metabolism community in liver cancer.** (A) Heatmap showing the level of DNA at the metabolism community CpGs in the TCGA-LIHC dataset (n=447). Red point indicates methylated CpGs and blue points indicate unmethylated CpGs. Rows represent CpGs and columns represent tumor samples. (B) Unsupervised hierarchical clustering of expression levels of genes in the metabolism community in liver cancer (n=377). Rows represent genes and columns represent tumor samples. (C) DNA methylation at the CpGs in the metabolism community in liver cancer (TCGA-LIHC). Box plot showing the expression of genes in the metabolism community in liver cancer (TCGA-LIHC). Wilcoxon-rank sum test p-values between tumor and normal samples are denoted in C and D. (E) Enrichment of the metabolism community CpGs in LIHC-defined regulatory regions obtained by Segway. The length of the bars represents -log10-transformed p-values obtained by hypergeometric testing using all the 450k probes as background. Only significant enrichments (BH-corrected p-value<0.05) are shown.

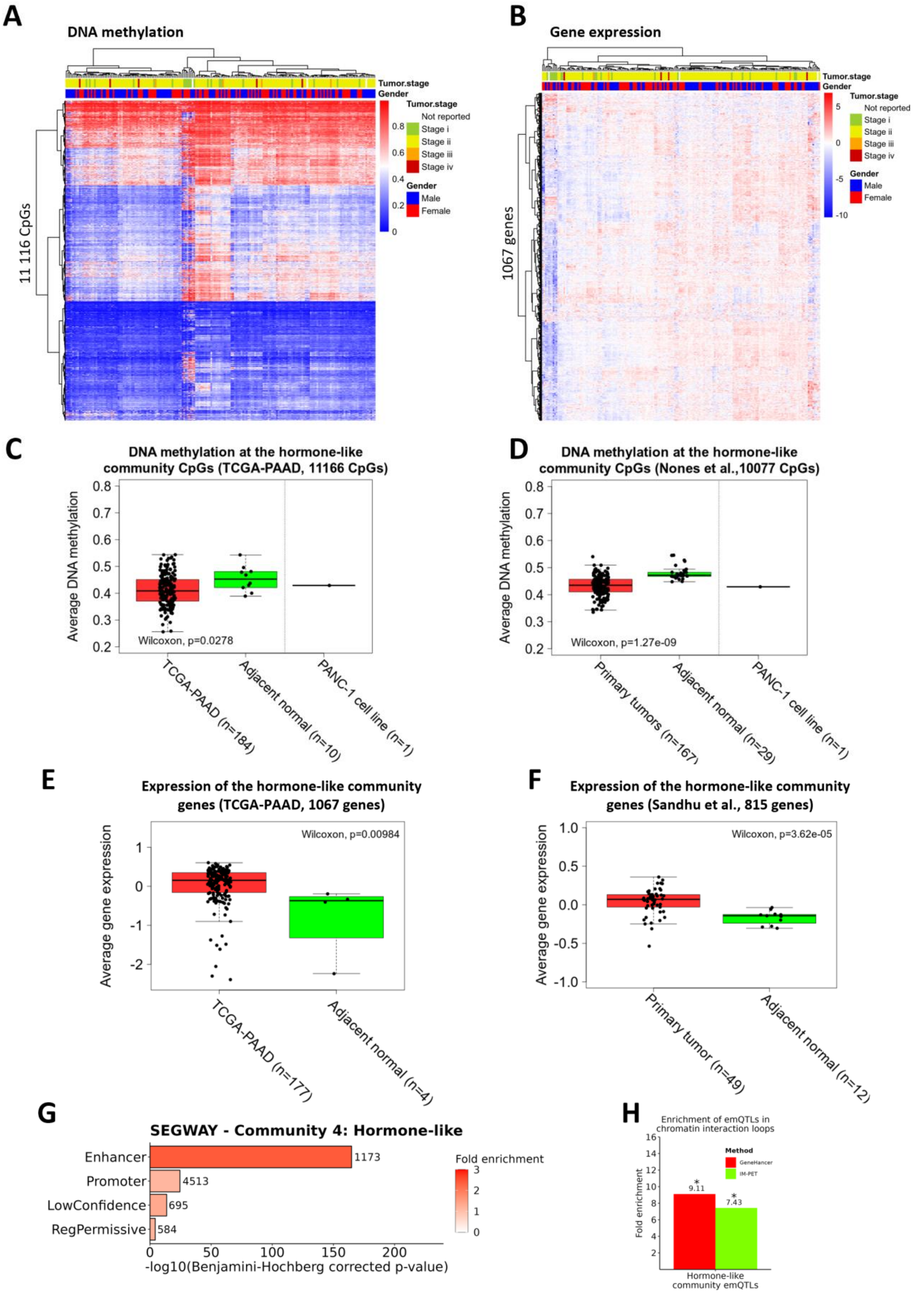


**Supplementary figure 17. emQTL community 4 is associated with hormone-related signaling.** (A) Unsupervised hierarchical clustering of the DNA methylation levels at the hormone-like community CpGs in pancreatic cancer (TCGA-PAAD). (B) Hierarchical clustering of the expression levels of the hormone-like community genes in pancreatic cancer (TCGA-PAAD). (E-F) Box plot showing the DNA methylation levels at the CpGs in the cell cycle community in TCGA (C) and the independent pancreatic cancer dataset obtained from Nones et al. [51] (D). DNA methylation levels for the PANC1 cell line are also included. Expression of the hormone-like community genes in pancreatic tumors from TCGA (E) and an independent cohort obtained from Sandhu et al. [54] (F). Wilcoxon-rank sum test p-values are denoted in each box plot. (G) Bar plot showing the enrichment of the hormone-like community CpGs in different regulatory regions of the genome according to SEGWAY from PANC1. (H) Enrichment of emQTL in the hormone-like community in IM-PET chromatin loops from the PANC1 pancreatic cancer cell line. The enrichment was determined using hypergeometric tests of significance using all possible *in cis* CpG-gene pairs as background. Significant enrichments after BH correction are marked with an asterisk.
